# Engineering potent avian GPR25 agonists using an ostrich *CXCL17* gene relic

**DOI:** 10.64898/2026.10.14.759250

**Authors:** Jie Yu, Hao-Zheng Li, Juan-Juan Wang, Ya-Li Liu, Zhan-Yun Guo

## Abstract

C-X-C motif chemokine ligand 17 (CXCL17) activates G protein-coupled receptor 25 (GPR25) in most vertebrate lineages. Although birds appear to have lost functional CXCL17, *CXCL17* gene relics and intact *GPR25* genes persist in certain taxa. This study investigated whether a fragment encoded by an avian *CXCL17* gene relic could guide the engineering of potent agonists for avian GPR25s. A 22-residue C-terminal fragment encoded by African ostrich *CXCL17* gene relic was grafted onto the corresponding position of loggerhead turtle or human CXCL17, and the resulting hybrids were prepared via bacterial overexpression and functionally characterized via cell-based assays. The turtle–ostrich and human–ostrich CXCL17 hybrids showed no detectable activity at turtle or human GPR25 but exhibited substantially greater potency than their parental proteins at two avian GPR25s in β-arrestin recruitment assays, with half-maximal effective concentrations of approximately 25 and 100 nM at the ostrich receptor and 5 and 13 nM at the mallard receptor, respectively. Both hybrids bound directly to the ostrich or mallard GPR25 in homogenous ligand−receptor binding assays and induced migration of transfected cells expressing the ostrich or mallard receptor in chemotaxis assays. These findings demonstrate that a fragment encoded by the ostrich *CXCL17* gene relic confers potent avian GPR25 agonism and establishes the C-terminus as the primary driver of species selectivity. These hybrids provide functional surrogate tools for probing avian GPR25 biology.

## Introduction

C-X-C motif chemokine ligand 17 (CXCL17) is a mucosal chemoattractant that recruits several leukocyte populations [1−9] and is implicated in the progression of multiple tumors [10−17]. Its chemoattractant activity is thought to be mediated by a cell surface receptor. However, the identity of this receptor has remained controversial for years, with the orphan receptor GPR35, the chemokine receptor CXCR4, and several MAS-related receptors proposed as candidates [18−23]. Recent independent studies, including our own, have demonstrated that CXCL17 activates the poorly characterized orphan G protein-coupled receptor 25 (GPR25) and this activation depends on its conserved C-terminal fragment [24−27], supporting GPR25 as a bona fide receptor for CXCL17.

GPR25 orthologs are widely distributed across vertebrates, from jawed fishes to mammals. By contrast, CXCL17 orthologs initially appeared to be restricted to mammals, raising questions about the evolutionary origin and phylogenetic distribution of the CXCL17−GPR25 signaling system. In recent studies, we identified and functionally characterized non-mammalian CXCL17 orthologs and paralogs in various jawed fishes, amphibians, and reptiles based on key amino acid sequence features and conserved gene synteny [28−32]. These findings suggest that the CXCL17−GPR25 signaling pair is evolutionarily conserved across most vertebrate lineages. Non-mammalian CXCL17 orthologs and paralogs share little overall sequence similarity with their mammalian counterparts, making them difficult to identify using conventional sequence homology-based approaches [28−32]. Before our studies, these non-mammalian CXCL17 orthologs or paralogs remained as uncharacterized proteins, were misannotated as non-coding transcripts, or lacked annotations in public databases [28−32]. Following our recent findings [28−32], public databases, including the National Center for Biotechnology Information (NCBI), the Zebrafish Information Network (ZFIN), and Xenbase, have updated annotations for several non-mammalian CXCL17 orthologs and paralogs.

Unlike other vertebrate lineages, birds appear to have lost CXCL17 during evolution, as we identified only nonfunctional *CXCL17* gene relics in some bird species [32]. These gene relics retain sequences encoding an N-terminal signal peptide and a C-terminal fragment with characteristic features of CXCL17 orthologs, including the essential Leu-Pro-Leu/Ile motif (Fig. S1). However, they contain frameshift or nonsense mutations, lack the typical exon−intron architecture, and show no evidence of expression. Consistent with the apparent loss of functional *CXCL17* gene, some neognathous birds, primarily members of the orders Galliformes, Passeriformes, and Psittaciformes, also appear to have lost the *GPR25* gene, according to the NCBI database [32]. However, intact *GPR25* genes remain in palaeognathous birds and some neognathous birds [32].

These observations raise two questions: do the identified avian *CXCL17* gene relics retain functionally relevant CXCL17 sequences, and can the remaining avian GPR25 receptors still be activated efficiently? To address these questions, we grafted a 22-residue C-terminal fragment encoded by the African ostrich (*Struthio camelus*) *CXCL17* gene relic onto the corresponding region of loggerhead turtle (*Caretta caretta*) CXCL17 (Cc-CXCL17) or human CXCL17 (Hs-CXCL17). Both hybrids exhibited substantially greater potency than their parental proteins at African ostrich GPR25 (Sc-GPR25) and mallard (*Anas platyrhynchos*) GPR25 (Ap-GPR25), representing palaeognathous and neognathous birds, respectively. However, neither hybrid showed detectable activity at loggerhead turtle GPR25 (Cc-GPR25) or human GPR25 (Hs-GPR25). Thus, this study provides two potent agonists for ostrich and mallard GPR25 and even some other avian GPR25s. The results support the functional relevance of the sequence retained in the ostrich *CXCL17* gene relic and demonstrate that the tested avian GPR25s remain capable of activation in heterologous cells. These findings also highlight the importance of the CXCL17 C-terminal fragment for species-dependent receptor selectivity, providing new insights into CXCL17 activating receptor GPR25.

## Materials and methods

### Amplification and sequencing of the ostrich *CXCL17* gene relic

Fresh liver of farmed African ostrich was purchased from market, and genomic DNA was extracted using an *EasyPure*^®^ Genomic DNA Kit (TranGen Biotech, Beijing, China). Thereafter, the ostrich *CXCL17* gene relic was PCR amplified using the extracted genomic DNA as template and synthetic oligoes as primers (forward: 5’-CGG GCT CCG GCC TCC GCC-3’; reverse: 5’-GGG GGC ATC CGA ATT CGG CCC-3’). The PCR product was analyzed by agarose gel electrophoresis and the band of approximately 300 bp was recovered from the gel by a *Trelief®* DNA Gel Extraction Kit (Tsingke Biotech, Beijing, China). Subsequently, the purified PCR product was either subjected to nanopore DNA sequencing (Tsingke Biotech, Beijing, China) or ligated into a pEASY vector using a *pEASY*^®^-Blunt Cloning Kit (TransGen). After the ligation product was transformed into Trans1-T1 phage resistant competent cells, plasmids were extracted from *Escherichia coli* colonies and subjected to Sanger DNA sequencing (Tsingke Biotech).

### Design and bacterial overexpression of CXCL17 hybrids

The bacterial expression constructs for human CXCL17 (Hs-CXCL17) and loggerhead turtle CXCL17 (Cc-CXCL17) were generated in our previous studies [32] and used as template for generating the coding region of CXCL17 hybrids (Fig. S2). In both hybrids, a 22-residue C-terminal fragment encoded by the ostrich *CXCL17* gene relic was grafted to the corresponding position of Cc-CXCL17 and Hs-CXCL17, respectively (Fig. 1A and Fig. S2). After three rounds of PCR amplification using pET/6×His-Cc-CXCL17 or pET/6×His-Hs-CXCL17 as template, the amplified DNA fragments were ligated into a pET expression vector via Gibson assembly, resulting in the expression constructs pET/Cc-Sc-CXCL17 and pET/Hs-Sc-CXCL17, respectively (Fig. S2). The C-terminally truncated or the N-terminally SmBiT-fused version of Cc-Sc-CXCL17 were also generated via PCR amplification and ligation into the pET vector via Gibson assembly (Fig. S2). The nucleotide sequence of these wild-type or mutant hybrids was confirmed by Sanger DNA sequencing (Tsingke Biotech).

**Fig. 1.**
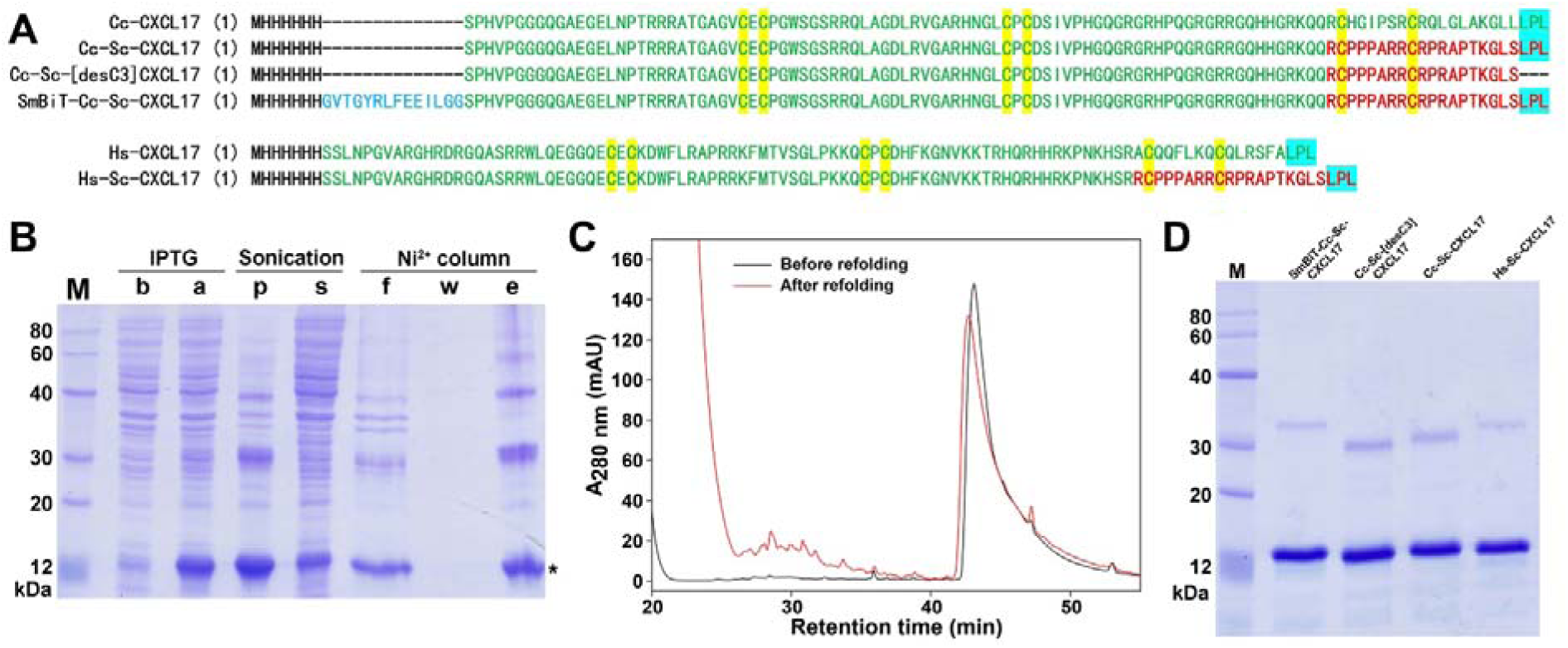
Preparation of recombinant CXCL17 hybrids. (**A**) Amino acid sequence of recombinant CXCL17s used in this study. The sequence of Hs-CXCL17 or Cc-CXCL17 is shown in green, that encoded by the ostrich *CXCL17* gene relic is shown in red. The SmBiT tag is shown in blue, and 6×His-tag in black. (**B**) SDS-PAGE analysis of Cc-Sc-CXCL17 samples at different preparation steps. Lane (M), protein ladder; lane (b), before IPTG induction; lane (a), after IPTG induction; lane (p), pellet after sonication; lane (s), supernatant after sonication; lane (f), flowthrough from the Ni^2+^ column; lane (w), washing fraction by 30 mM imidazole; lane (e), eluted fraction by 250 mM imidazole. After electrophoresis, the gel was stained by Coomassie brilliant blue R250. Band of the monomeric hybrid is indicated by an asterisk. (**C**) HPLC analysis of the recombinant Cc-Sc-CXCL17 before or after *in vitro* refolding. (**D**) SDS-PAGE analysis of the recombinant CXCL17 hybrids after *in vitro* refolding and HPLC purification. The gel was stained by Coomassie brilliant blue R250 after electrophoresis.

Thereafter, CXCL17 hybrids were prepared according to our previous procedures developed for other CXCL17s [28−32]. Briefly, the N-terminally 6×His-tagged hybrids were overexpressed in the *E. coli* strain BL21(DE3) as inclusion bodies, solubilized via an *S*-sulfonation approach, and purified by Ni^2+^ affinity chromatography. After *in vitro* refolding, the hybrids were further purified by HPLC using a C_18_ reverse-phase column. The eluted fraction was manually collected, lyophilized, dissolved in 1.0 mM aqueous hydrochloride (pH 3.0), and quantified by ultra-violet absorbance at 280 nm according to their theoretical extinction coefficient (5875 M^-1^ cm^-1^ for full-length or C-terminally truncated Cc-Sc-CXCL17; 7365 M^-1^ cm^-1^ for the SmBiT-fused Cc-Sc-CXCL17; 11375 M^-1^ cm^-1^ for Hs-Sc-CXCL17). During purification, samples at different preparation steps were analyzed by SDS-PAGE.

### Generation of expression constructs for ostrich GPR25

The expression constructs for the mallard GPR25 (Ap-GPR25) were generated in our previous study via chemical synthesis of its coding region according to the NCBI reference sequence and subsequent cloning into appropriate expression vectors [32]. The coding region of the African ostrich GPR25 (Sc-GPR25) was PCR amplified using the extracted ostrich genomic DNA as template and synthetic oligos as primers (Table S1). The amplified DNA fragment of ∼1.2 kb was recovered from agarose gel after electrophoresis and ligated into a pEASY vector using a *pEASY*^®^-Blunt Cloning Kit (TransGen) (Table S1). After plasmid was extracted from transformed *E. coli* colonies, Sanger DNA sequencing confirmed its nucleotide sequence is identical to the reference sequence XM_009674745 (Fig. S3). Subsequently, the coding region of Sc-GPR25 was PCR amplified using the pEASY/Sc-GPR25 as template and ligated into appropriate Dox-inducible expression vectors (Table S1). The construct pTRE3G-BI/Sc-GPR25-LgBiT:SmBiT-ARRB2 coexpresses a C-terminally LgBiT-fused Sc-GPR25 (Sc-GPR25-LgBiT) and an N-terminally SmBiT-fused human β-arrestin 2 (SmBiT-ARRB2) for NanoBiT-based β-arrestin recruitment assays (Fig. S3). The expression construct PB-TRE/sLgBiT-Sc-GPR25 expresses an N-terminally sLgBiT-fused Sc-GPR25 (sLgBiT-Sc-GPR25) for NanoBiT-based homogenous binding assays, while the construct PB-TRE/Sc-GPR25 expresses an untagged receptor for chemotaxis assays (Fig. S3). The nucleotide sequence of Sc-GPR25 in these expression constructs was confirmed by Sanger DNA sequencing (Tsingke Biotech).

### The NanoBiT-based β-arrestin recruitment assays

The NanoBiT-based β-arrestin recruitment assays for avian GPR25s were conducted according to our previous procedure developed for other receptors [28−32]. Briefly, HEK293T cells were transiently cotransfected with the β-arrestin recruitment assay construct and the expression control plasmid pCMV-Tet3G (Clontech, Mountain View, CA, USA) using transfection reagent Lipo8000 (Beyotime Biotech, Shanghai, China). Next day, the transfected cells were trypsinized, seeded into white opaque 96-well plates, and cultured in induction medium (complete DMEM plus 1.0 ng/mL of Dox) for ∼24 h to ∼90% confluence. Thereafter, the induction medium was removed, NanoLuc substrate-containing activation solution (serum-free DMEM plus 1% BSA) was added (40 μL/well, containing 1.0 μL of NanoLuc substrate stock from Promega, Madison, WI, USA), and bioluminescence data were collected on a SpectraMax iD3 plate reader (Molecular Devices, Sunnyvale, CA, USA) for ∼4 min. Subsequently, indicated CXCL17 protein was added (10 μL/well, diluted in activation solution) and bioluminescence data were continuously collected for ∼10 min. The measured bioluminescence data were corrected for inter well variability by forcing all curves after addition of NanoLuc substrate to same level and plotted using the SigmaPlot 10.0 software (SYSTAT software, Chicago, IL, USA). To generate the dose-response curve, the highest bioluminescence data were plotted with the corresponding agonist concentrations using the SigmaPlot 10.0 software and EC_50_ was calculated from the curve by the software (SYSTAT software).

### The NanoBiT-based ligand−receptor binding assays

The NanoBiT-based homogenous binding assays for avian GPR25s were conducted according to our previous procedure using recombinant SmBiT-Cc-Sc-CXCL17 as a binding tracer [28−32]. Briefly, HEK293T cells were cotransfected with the NanoBiT-based binding assay construct and the tyrosylprotein sulfotransferase expression construct pTRE3G-BI/TPST1:TPST2. Next day, the transfected cells were trypsinized, seeded into white opaque 96-well plates, and cultured in induction medium (complete DMEM plus 20 ng/mL of Dox) for ∼24 h to ∼90% confluence. Thereafter, the induction medium was removed and pre-warmed binding solution (serum-free DMEM plus 0.1% BSA and 0.01% Tween-20) was added (50 μL/well). The binding solution either contained varied concentrations of the binding tracer (for saturation binding) or contained a constant concentration of the binding tracer and varied concentrations of a competitor (for competition binding). After incubation at room temperature for ∼1 h, diluted NanoLuc substrate (30-fold dilution in the binding solution) was added (10 μL/well), and bioluminescence data were immediately measured on a SpectraMax iD3 plate reader (Molecular Devices). The measured bioluminescence data were expressed as mean ± SD (*n* = 3) and fitted to one-site binding model using the SigmaPlot 10.0 software (SYSTAT software).

### Chemotaxis assays of transfected HEK293T cells expressing avian GPR25

The chemotaxis assays of the transfected HEK293T cells were conducted using permeable transwell inserts coated with polyethylene terephthalate membrane (8 μm pore size) according to our previous procedure [28−32]. Briefly, transiently transfected HEK293T cells were cultured in induction medium (complete DMEM plus 1.0 ng/mL of Dox) for ∼24 h to induce expression of untagged avian GPR25. Thereafter, the cells were trypsinized, suspended in serum-free DMEM (∼5×10^5^ cells/mL), and seeded into the transwell inserts (200 μL/well). The inserts were put into a 24-well plate containing indicated CXCL17 proteins (diluted in serum-free DMEM plus 0.2% BSA, 500 μL/well) and incubated at 37°C for ∼5 h. Thereafter, solution in the inserts was removed and cells on the upper face of the permeable membrane were wiped off using cotton swaps, and cells on the lower face of the permeable membrane were fixed with 4% paraformaldehyde solution, stained with crystal violet staining solution (Beyotime Technology), and observed under a microscope. The migrated cells were quantitated using the ImageJ software and the results were expressed as mean ± SD (*n* = 3).

## Results

### Confirmation of the *CXCL17* gene relic in African ostrich

In the African ostrich reference genome bStrCam1.hap1 (NC_090980.1) at NCBI gene database, a *CXCL17* gene relic lies adjacent to the *LIPE* gene on chromosome 39 (Fig. S4). This relic is only 334 bp long and retains sequences encoding a typical N-terminal signal peptide and a C-terminal peptide with characteristic features of CXCL17 orthologs, including the essential Leu-Pro-Leu motif (Fig. S5A). However, it contains a frameshift mutation, and the RNA sequencing data available in the NCBI gene database show no evidence of transcription (Fig. S4). Functional *CXCL17* genes in mammals, reptiles, amphibians, and fishes typically comprise four exons and three introns and span thousands of base pairs. By contrast, all avian *CXCL17* gene relics identified to date span only 300−400 bp and contain frameshift or nonsense mutations [32], suggesting that they can no longer encode functional CXCL17 proteins.

To verify the presence of the *CXCL17* gene relic in the African ostrich, we extracted genomic DNA from fresh liver tissue obtained from a farmed ostrich and amplified the region by PCR. An approximately 300 bp fragment was obtained, and DNA sequencing confirmed that it corresponded to the expected *CXCL17* gene relic (Fig. S5B). The amplified sequence differed slightly from the reference sequence, with three indels and one nucleotide substitution (Fig. S5C). Nevertheless, both sequences encoded the same C-terminal fragment with characteristic features of CXCL17 orthologs. These results confirm the presence of the *CXCL17* gene relic in the African ostrich genome and suggest possible sequence polymorphism among individuals.

### Design and preparation of recombinant CXCL17 hybrids

To assess the functional relevance of the sequence retained in the African ostrich *CXCL17* gene relic, we grafted its encoded C-terminal 22 residues onto the corresponding region of Cc-CXCL17 and Hs-CXCL17. This design was based on the importance of the C-terminal fragment for the function of CXCL17 orthologs and paralogs [24−32]. The resulting hybrids, Cc-Sc-CXCL17 and Hs-Sc-CXCL17, each contain the short C-terminal fragment encoded by the ostrich *CXCL17* gene relic (red) and the longer N-terminal region of Cc-CXCL17 or Hs-CXCL17 (green; Fig. 1A). Both hybrids carry an N-terminal 6×His tag to facilitate purification.

Following IPTG induction in *E. coli*, Cc-Sc-CXCL17 produced a prominent band (indicated by an asterisk) at approximately 12 kDa on SDS-PAGE, consistent with overexpression of the hybrid (Fig. 1B). The protein was solubilized from inclusion bodies by *S*-sulfonation and purified using a Ni^2+^ affinity column. SDS-PAGE revealed both monomers and oligomers (Fig. 1B), suggesting that Cc-Sc-CXCL17 is prone to intermolecular cross-linking, as observed for other CXCL17 proteins [28−32]. After *in vitro* refolding, Cc-Sc-CXCL17 was further purified by HPLC, yielding a major elution peak from a C_18_ reverse-phase column (Fig. 1C). Following HPLC purification and lyophilization, the refolded protein showed a prominent monomer band and a faint dimer band on SDS-PAGE (Fig. 1D). The other CXCL17 hybrids were prepared using the same procedure and also showed predominantly monomeric bands (Fig. 1D). These results demonstrate the successful production of recombinant CXCL17 hybrids through bacterial expression, refolding, and purification.

### Activation of avian GPR25s by recombinant CXCL17 hybrids

We assessed the activity of the recombinant CXCL17 hybrids using the NanoBiT-based β-arrestin recruitment assay, which uses a C-terminally LgBiT-fused receptor coexpressed with SmBiT-ARRB2. Upon activation, the receptor recruits SmBiT-ARRB2, bringing SmBiT and LgBiT into proximity. Complementation of the two fragments generates bioluminescence, allowing receptor activation to be monitored in real time.

Addition of Cc-CXCL17 to live HEK293T cells coexpressing Sc-GPR25-LgBiT and SmBiT-ARRB2 increased bioluminescence in a concentration-dependent manner (Fig. 2A). However, its EC_50_ could not be reliably determined within the tested concentration range (Fig. 2A, inset), indicating low cross-species potency at Sc-GPR25. By contrast, Cc-Sc-CXCL17 exhibited substantially greater potency, with an EC_50_ of approximately 25 nM (Fig. 2B). This result indicates that the C-terminal fragment encoded by the African ostrich *CXCL17* gene relic markedly enhances activation of Sc-GPR25. The C-terminally truncated mutant Cc-Sc-[desC3]CXCL17 showed no detectable activity at Sc-GPR25 (Fig. 2C), indicating that the terminal Xaa-Pro-Yaa motif is essential for the hybrid activity, consistent with findings for other CXCL17 orthologs and paralogs [28−32].

**Fig. 2.**
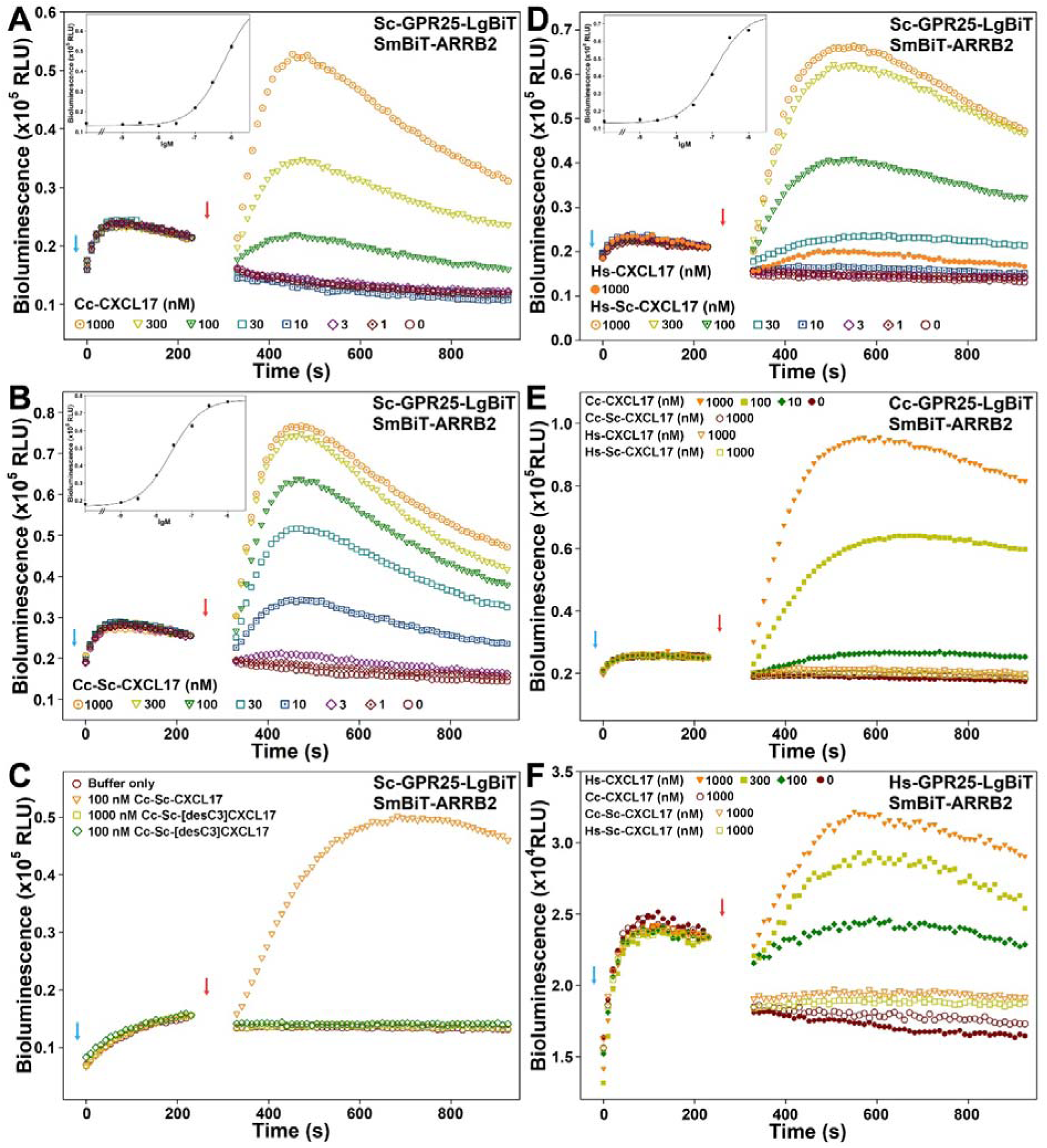
NanoBiT-based β-arrestin recruitment assays for Sc-GPR25, Cc-GPR25, and Hs-GPR25. (**A−D**) Typical bioluminescence change upon stimulation of Sc-GPR25 by Cc-CXCL17 (A), Cc-Sc-CXCL17 (B), Cc-Sc-[desC3]CXCL17 (C), or Hs-Sc-CXCL17 and Hs-CXCL17 (D). Inner panels, dose-response curve of Cc-CXCL17 (A), Cc-Sc-CXCL17 (B), or Hs-Sc-CXCL17 (D). (**E,F**) Typical bioluminescence change upon stimulation of Cc-GPR25 (E) or Hs-GPR25 (F) by CXCL17 hybrids, Cc-CXCL17, and Hs-CXCL17. In these NanoBiT-based β-arrestin recruitment assays, NanoLuc substrate and indicated recombinant CXCL17 protein were sequentially added to living HEK293T cells coexpressing the indicated C-terminally LgBiT-fused receptor and SmBiT-ARRB2, and bioluminescence was monitored in a real-time manner on a plate reader. To obtain the dose-response curve, the highest bioluminescence upon agonist stimulation was plotted versus agonist concentrations via SigmaPlot10.0 software and EC_50_ values were calculated from the sigmoidal curves by the software. Blue arrows indicate the addition of NanoLuc substrate, and red arrows indicate the addition of CXCL17 protein.

Hs-CXCL17 showed very little cross-species activity at Sc-GPR25 in the β-arrestin recruitment assay (Fig. 2D). In comparison, Hs-Sc-CXCL17 exhibited substantially greater potency, with an EC_50_ of approximately 100 nM (Fig. 2D), further supporting the importance of the grafted C-terminal fragment for activation of the ostrich receptor. At Cc-GPR25, Cc-CXCL17 induced a rapid, concentration-dependent increase in bioluminescence, whereas neither hybrids nor Hs-CXCL17 showed detectable activity (Fig. 2E). Both hybrids and Cc-CXCL17 also showed negligible activity at Hs-GPR25 (Fig. 2F). These findings highlight the importance of the CXCL17 C-terminal fragment for species-dependent receptor selectivity.

Cc-CXCL17 also activated Ap-GPR25 (Fig. 3A), with an EC_50_ of approximately 120 nM (Fig. 3D). However, the bioluminescence response was much smaller than that observed for Sc-GPR25, making receptor activation more difficult to quantify. Cc-Sc-CXCL17 exhibited substantially greater potency than Cc-CXCL17 (Fig. 3B), with an EC_50_ of approximately 5 nM and a moderately higher E (Fig. 3D). Removal of the three C-terminal residues abolished activity at Ap-GPR25 (Fig. 3B), confirming the importance of the terminal Xaa-Pro-Yaa motif for activation of the mallard receptor. Hs-CXCL17 showed minimal activity at Ap-GPR25 even at 1000 nM (Fig. 3C), whereas Hs-Sc-CXCL17 exhibited substantially greater potency (Fig. 3C), with an EC_50_ of approximately 13 nM (Fig. 3D). These results demonstrate that the C-terminal fragment encoded by the African ostrich *CXCL17* gene relic is a key determinant of hybrid activity at avian GPR25s.

**Fig. 3.**
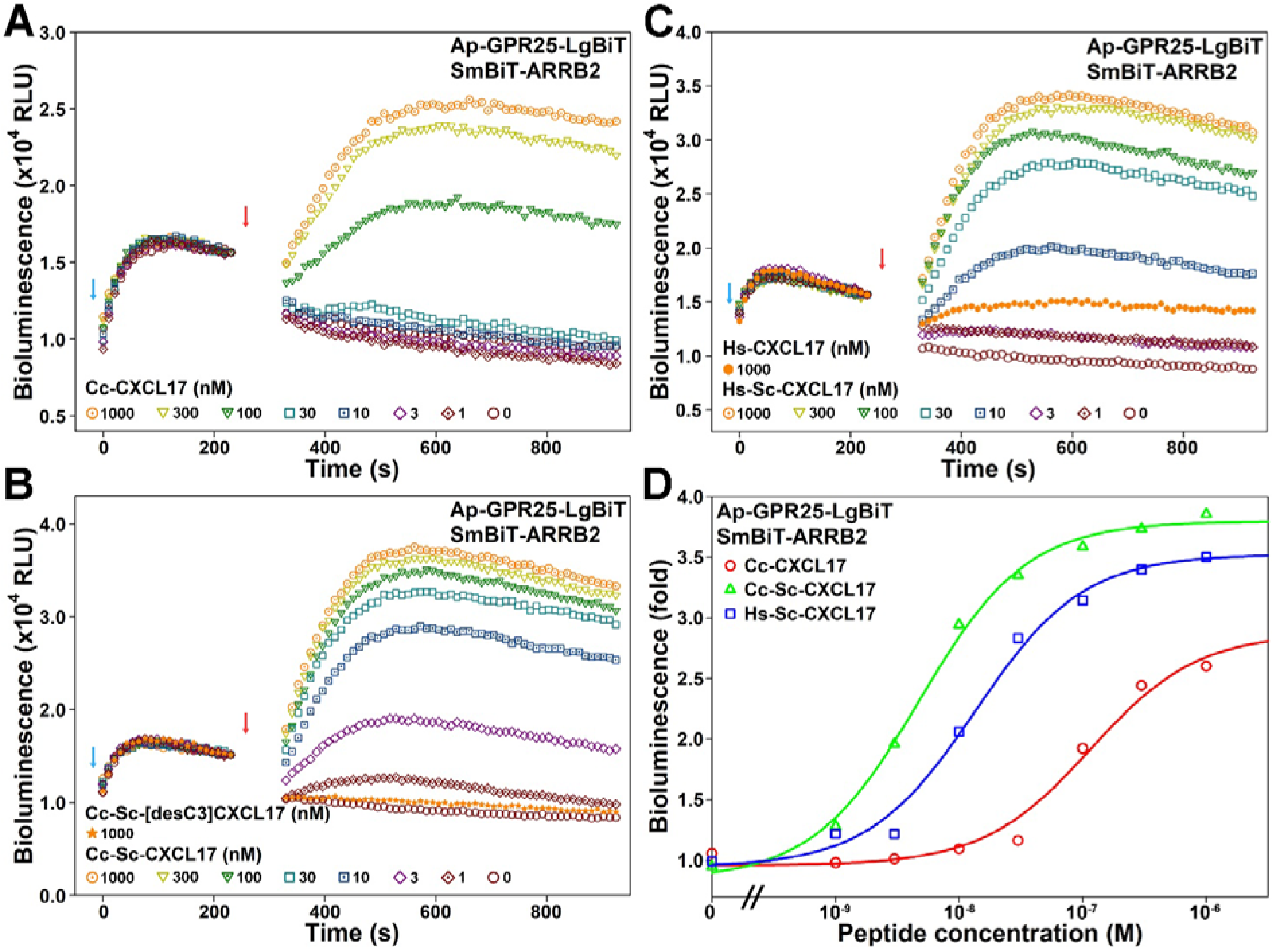
NanoBiT-based β-arrstin recruitment assays for Ap-GPR25. (**A−C**) Typical bioluminescence change upon stimulation of Ap-GPR25 by Cc-CXCL17 (A), the full-length or C-terminally truncated Cc-Sc-CXCL17 (B), or Hs-CXCL17 and Hs-Sc-CXCL17 (C). (**D**) Dose-response curves of Cc-CXCL17, Cc-Sc-CXCL17, and Hs-Sc-CXCL17 at Ap-GPR25 measured by the β-arrstin recruitment assays. In these assays, NanoLuc substrate and indicated CXCL17 protein were sequentially added to living HEK293T cells coexpressing Ap-GPR25-LgBiT and SmBiT-ARRB2, and bioluminescence was monitored in a real-time manner on a plate reader. To obtain the dose-response curve in panel D, the highest bioluminescence upon agonist stimulation in panel A−C was plotted versus agonist concentrations via SigmaPlot10.0 software and EC_50_ values were calculated from the sigmoidal curves by the software. Blue arrows the indicate addition of NanoLuc substrate, and red arrows indicate the addition of CXCL17 protein.

### Direct binding of recombinant CXCL17 hybrids to avian GPR25s

We assessed direct binding of the CXCL17 hybrids to avian GPR25s using the NanoBiT-based homogeneous binding assay, which uses an N-terminally sLgBiT-fused receptor and a SmBiT-fused ligand as a tracer. Following recombinant expression and purification, the tracer SmBiT-Cc-Sc-CXCL17 retained the ability to activate Sc-GPR25 in the NanoBiT-based β-arrestin recruitment assay, although its activity was lower than that of Cc-Sc-CXCL17 (Fig. S6), supporting its use as a tracer in the binding assay.

Addition of SmBiT-Cc-Sc-CXCL17 to live HEK293T cells overexpressing sLgBiT-Sc-GPR25 yielded a saturation binding curve (Fig. 4A), with a calculated *K*_d_ value of 37.6 ± 8.6 nM (*n* = 3). Competition with 1.0 μM Cc-Sc-CXCL17 markedly reduced bioluminescence (Fig. 4A), confirming specific binding of the tracer to Sc-GPR25. In competition binding assays, Cc-Sc-CXCL17 displaced the tracer in a concentration-dependent manner (Fig. 4B), producing a sigmoidal inhibition curve with an IC_50_ of 2.4 ± 0.4 nM (*n* = 3). Hs-Sc-CXCL17 also displaced the tracer, but its IC_50_ was higher, at 17.8 ± 2.7 nM (Fig. 4B), consistent with its lower potency in the β-arrestin recruitment assay. Both parental proteins, Cc-CXCL17 and Hs-CXCL17, exhibited substantially lower binding potency than the hybrids, with IC_50_ values of approximately 600 nM (Fig. 4B). The C-terminally truncated mutant Cc-Sc-[desC3]CXCL17 showed no detectable binding to Sc-GPR25 (Fig. 4B), consistent with its lack of activity in the β-arrestin recruitment assay.

**Fig. 4.**
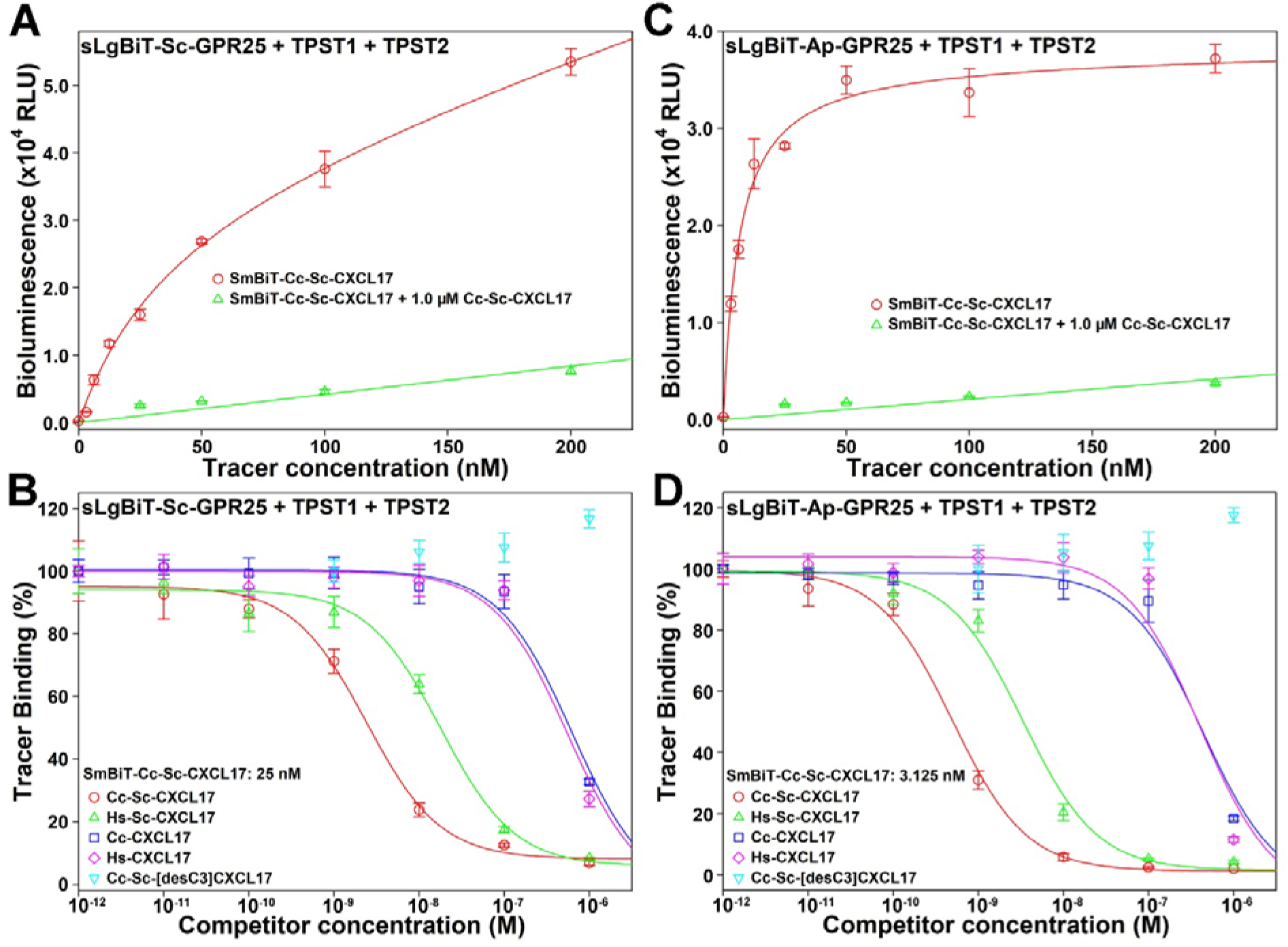
NanoBiT-based homogenous ligand-binding assays of Sc-GPR25 and Ap-GPR25. (**A,C**) Saturation binding of SmBiT-Cc-Sc-CXCL17 with sLgBiT-Sc-GPR25 (A) or sLgBiT-Ap-GPR25 (C). (**B,D**) Competition binding of various CXCL17s with sLgBiT-Sc-GPR25 (B) or sLgBiT-Ap-GPR25 (D) using SmBiT-Cc-Sc-CXCL17 as a tracer. In these binding assays, living HEK293T cells coexpressing the N-terminally sLgBiT-fused GPR25 and human tyrosylprotein sulfotransferases (TPST1 and TPST2) were incubated with the SmBiT-Cc-Sc-CXCL17 tracer under absence or presence of a competitor for 1 h. Thereafter, diluted NanoLuc substrate was added and bioluminescence was measured on a plate reader. The measured bioluminescence data were expressed as mean ± SD (*n* = 3) and fitted to one-site binding model using SigmaPlot10.0 software. K_d_ values were calculated from the saturation binding curves and IC_50_ values were calculated from the competition binding curves by the software.

The tracer also exhibited a typical hyperbolic saturation binding curve at sLgBiT-Ap-GPR25 (Fig. 4C), with a *K*_d_ value of 6.5 ± 0.9 nM (*n* = 3), indicating high-affinity binding to the mallard receptor. In competition binding assays, both hybrids showed high binding potency at Ap-GPR25, with IC_50_ values of 0.50 ± 0.05 nM for Cc-Sc-CXCL17 and 3.3 ± 0.5 nM for Hs-Sc-CXCL17 (Fig. 4D). By contrast, both parental proteins had substantially lower binding potency, with IC_50_ values of approximately 400 nM (Fig. 4D). The C-terminally truncated mutant again showed no detectable binding to Ap-GPR25 (Fig. 4D), confirming the importance of the terminal Xaa-Pro-Yaa motif for CXCL17 binding to avian GPR25s.

### Chemotactic migration of HEK293T cells expressing avian GPR25

We evaluated the cellular response to avian GPR25 activation using chemotaxis assays (Fig. 5). Both Cc-Sc-CXCL17 and Hs-Sc-CXCL17 induced concentration-dependent migration of transfected HEK293T cells expressing Sc-GPR25, with a modest but significant effect at 10 nM (Fig. 5A,B). By contrast, the C-terminally truncated mutant induced only minimal migration, even at 1000 nM (Fig. 5A,B). Both CXCL17 hybrids also induced chemotactic migration of HEK293T cells expressing Ap-GPR25 (Fig. 5C,D). In non-transfected HEK293T cells, however, both hybrids had minimal effects at concentrations up to 1000 nM (Fig. S7). These results demonstrate that recombinant CXCL17 hybrids can induce chemotactic migration of cells expressing ostrich or mallard GPR25, supporting the functional capacity of these avian receptors in a heterologous cell system.

**Fig. 5.**
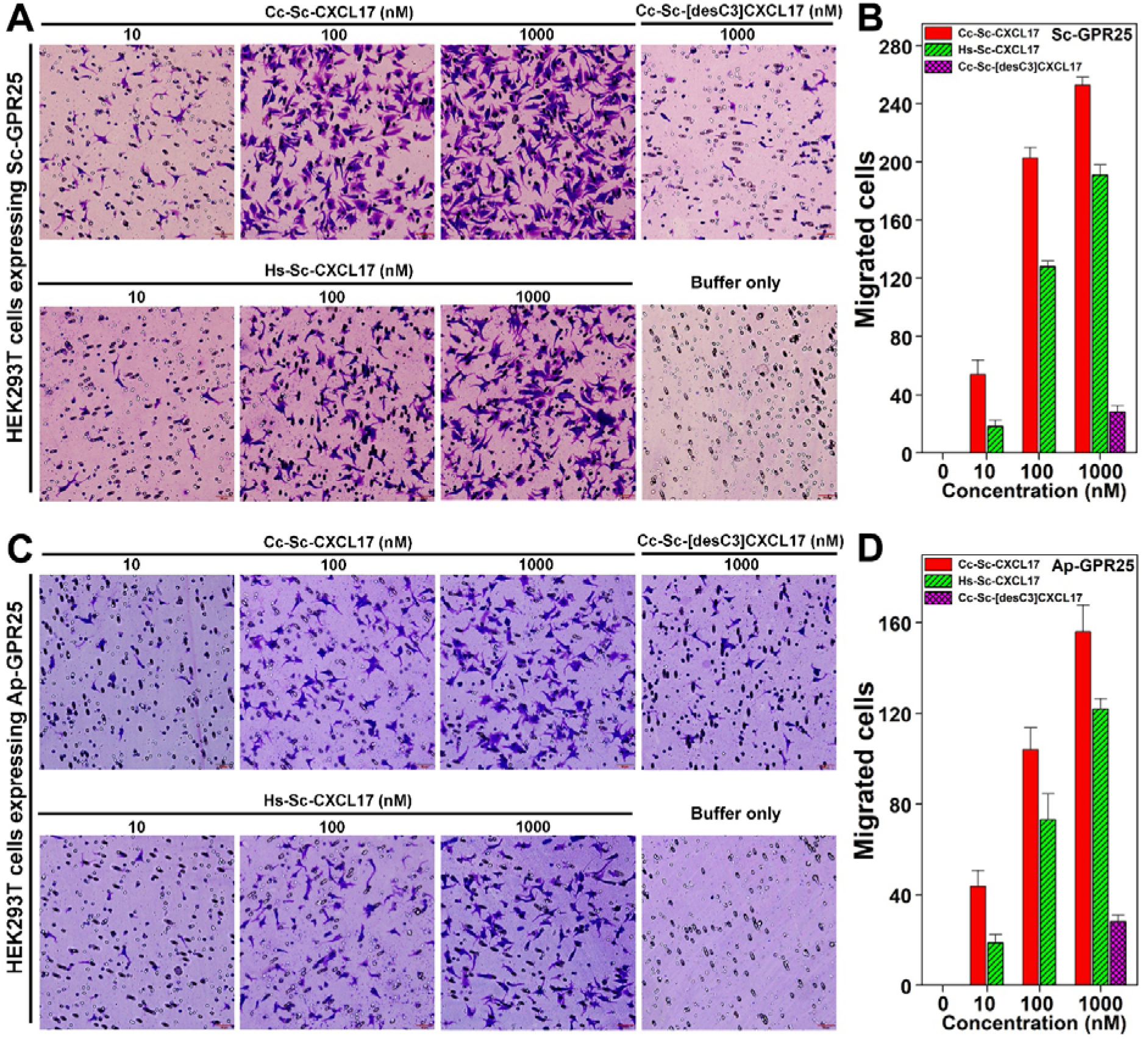
Chemotaxis assays of transfected HEK293T cells expressing Sc-GPR25 or Ap-GPR25. (**A,C**) Representative images of the migrated HEK293T cells expressing Sc-GPR25 (A) or Ap-GPR25 (C) upon induction by full-length or C-terminally truncated CXCL17 hybrid. The scale bar in these images is 50 μm. (**B,D**) Quantitative analysis of the migrated HEK293T cells expressing Sc-GPR25 (B) or Ap-GPR25 (D) upon induction by CXCL17 hybrids. In these transwell assays, transfected HEK293T cells were seeded into permeable membrane-coated inserts and induced by chemotactic solution containing indicated CXCL17 hybrid. After being cultured at 37°C for ∼5 h, cells on the upper face of the permeable membrane were wiped off, and cells on the lower face were fixed, stained, and observed under a microscope. Representative images are shown in panel A and C, the cell numbers in three images were analyzed using the ImageJ software, expressed as mean ± SD (*n* = 3), and shown in panel B and D.

## Discussion

In this study, we developed two potent agonists for ostrich and mallard GPR25s by grafting a C-terminal fragment encoded by the ostrich *CXCL17* gene relic onto the corresponding region of loggerhead turtle or human CXCL17. These findings support the functional relevance of the CXCL17 sequence retained in the ostrich *CXCL17* gene relic. Given the high sequence similarity among the remaining avian GPR25 orthologs, the two hybrids may also activate receptors from other bird species [32]. The same strategy could be used to develop additional avian GPR25 agonists by incorporating C-terminal fragments encoded by other avian *CXCL17* gene relics into CXCL17 orthologs from other vertebrate lineages. Our study therefore establishes a strategy for engineering potent avian GPR25 agonists. Such agonists provide valuable tools for investigating receptor function, particularly because birds appear to have lost the functional *CXCL17* gene during evolution [32]. Our results also identify the CXCL17 C-terminal fragment as a key determinant of species-dependent receptor selectivity, providing new insights into the molecular basis of CXCL17 recognition by GPR25.

Activation of ostrich and mallard GPR25s by the engineered CXCL17 hybrids demonstrates that these receptors retain functional capacity in heterologous cells. However, whether avian GPR25 is expressed *in vivo* remains largely unknown. Available human data indicate that endogenous *GPR25* expression is very low. According to the Human Protein Atlas, the highest reported *GPR25* transcript levels are 1.8 nTPM across tissues and 2.3 nCPM across single-cell populations (https://www.proteinatlas.org/ENSG00000170128-GPR25). By contrast, the highest reported *CXCL17* transcript levels are 338 nTPM in the stomach and 2922 nCPM in apical squamous epithelial cells of the esophagus (https://www.proteinatlas.org/search/CXCL17). Furthermore, because *GPR25* is an intronless gene, genomic DNA contamination can confound measurements of its transcript abundance. Establishing the endogenous expression patterns of avian *GPR25* will therefore be essential for assessing its potential physiological roles in birds.

Despite the apparent loss of the functional *CXCL17* gene in the avian lineage, palaeognathous birds and some neognathous birds retain intact *GPR25* genes [32]. The physiological roles of these receptors in the apparent absence of functional CXCL17 remain unclear. Our recent findings suggest that apelin and apela, two peptide hormones highly conserved across vertebrates, are unlikely to act as avian GPR25 agonists, despite the close evolutionary relationship between avian GPR25 and avian apelin receptors (APLNRs) [32]. Further studies are therefore needed to determine whether the retained avian GPR25 receptors have physiological functions and whether they recognize alternative endogenous agonists.

## Data availability statement

The data of this study are available in this manuscript, as well as the associated supplementary information.

## CRediT Author Contribution

Conceptualization, Z.Y.G.; investigation, J.Y., H.Z.L., and J.J.W.; writing – original draft, Z.Y.G.; writing – review & editing, Z.Y.G., and J.Y.; formal analysis, Y.L.L.; project administration, Y.L.L.; funding acquisition, Z.Y.G.; supervision, Z.Y.G. All authors read and approved the final manuscript.

## Abbreviations

BSA: bovine serum albumin;
CXCL17: C-X-C motif chemokine ligand 17
DMEM: Dulbecco’s modified Eagle medium
Dox: doxycycline
EC_50_: half-maximal effective concentration
E_max_: maximal effect
GPR25: G protein-coupled receptor 25
HEK: human embryonic kidney
HPLC: high performance liquid chromatography
IC_50_: half-maximal inhibitory concentration
IPTG: isopropyl-β-D-thiogalactopyranoside
LgBiT: large NanoLuc fragment for NanoBiT
NanoBiT: NanoLuc Binary Technology
NanoLuc: nanoluciferase
NCBI: National Center for Biotechnology Information
nCPM: normalized counts per million
nTPM: normalized transcripts per million
PCR: polymerase chain reaction
SD: standard deviation
SDS-PAGE: sodium dodecyl sulfate-polyacrylamide gel electrophoresis
sLgBiT: secretory LgBiT
SmBiT: small NanoLuc fragment for NanoBiT.

