## Supplementary material for "Engineering potent avian GPR25 agonists using an ostrich *CXCL17* gene relic": Table S1 and Fig. S1-S7


**Contents**


**Table S1.** Information for generation of the expression constructs for Sc-GPR25.

**Fig. S1.** Partial amino acid sequences encoded by the newly identified avian *CXCL17* gene relics.

**Fig. S2.** The nucleotide sequence and amino acid sequence of hybrid CXCL17s carrying a C-terminal fragment encoded by African ostrich *CXCL17* gene relic.

**Fig. S3.** The nucleotide sequence and amino acid sequence of Sc-GPR25 in various expression constructs.

**Fig. S4.** Position of the *CXCL17* gene relic in reference genome of African ostrich (*Struthio camelus*).

**Fig. S5.** Confirming presence of *CXCL17* gene relic in African ostrich.

**Fig. S6.** Activity of the tracer SmBiT-Cc-Sc-CXCL17 measured via the NanoBiT-based β-arrestin recruitment assay.

**Fig. S7.** Presentative images of migrated non-transfected HEK293T cells after induced by Cc-Sc-CXCL17 or Hs-Sc-CXCL17 in transwell chemotaxis assays.

**Table S1.** Information for generation of the expression constructs for Sc-GPR25.

| Expression constructs | Vectors for cloning | Restriction enzymes cleaving the vector | Primers for PCR amplification (5' to 3') | Template for PCR amplification | Approach for construct generation |
|---|---|---|---|---|---|
| pEASY/ Sc-GPR25 | pEASY-Blunt cloning vector | no | Forward: GGC GCT GGG GAT GCC GAC<br>Reverse: CAA GCT CAG CCC TCC CCA AAT | Genomic DNA extract from liver of African ostrich | Blunt end ligation |
| pTRE3G-BI/ Sc-GPR25-LgBiT: SmBiT-ARRB2 | pTRE3G-BI/ GPR182-LgBiT: SmBiT-ARRB2 | NheI; AgeI (removing GPR182) | Forward: TT CGG GGA GAC CCA AGC TGG ATG GGT CCC CAA GCA GCC GA<br>Reverse: GCT AGA CCC TCC GCC GGT ACC GAC CGG TGG CTC GTT TGG CCC CCA ATC CT | pEASY/ Sc-GPR25 | Gibson assembly |
| PB-TRE/ sLgBiT-Sc-GPR25 | PB-TRE/ sLgBiT-GPR182 | KpnI; PmeI (removing GPR182) | Forward: GGT GGC AGC GGC GGT GGT ACC GGT CCC CAA GCA GCC GAG GA<br>Reverse: GAG GCT GAT CAG CGG GTT T CTA CTC GTT TGG CCC CCA | pEASY/ Sc-GPR25 | Gibson assembly |
| PB-TRE/ Sc-GPR25 | PB-TRE/ dCas9-VPR | NheI; PmeI (removing dCas9-VPR) | Forward: CCT CGT AAA GGT CTA GAG ATG GGT CCC CAA GCA GCC<br>Reverse: AGT TAG CCT CCC CCG TTT CTA CTC GTT TGG CCC CCA | pEASY/ Sc-GPR25 | Gibson assembly |

PB-TRE/sLgBiT-GPR182 was generated in our laboratory based on PB-TRE/dCas9-VPR (Addgene cat#: 63800) via removal of the dCas9-VPR fragment by NheI and PmeI cleavage and then ligation of sLgBiT-GPR182 fragment, unpublished data.

pTRE3G-BI/GPR182-LgBiT:SmBiT-ARRB2 was generated in our laboratory, unpublished data.

For oligo primers, the sequence pairing with vector in Gibson assembly is highlighted in yellow, the sequence pairing with PCR template is underlined.

**A**

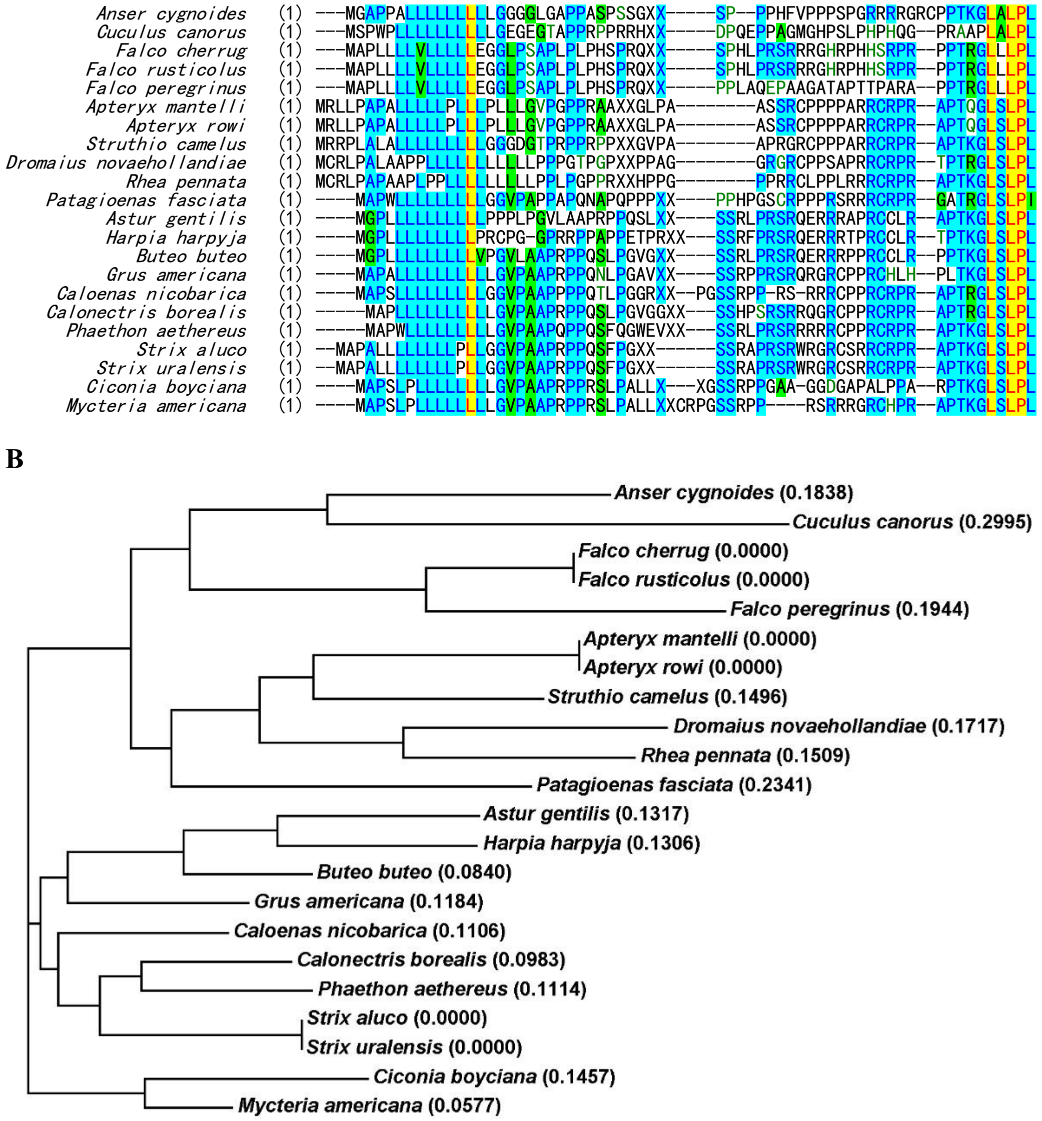

**Fig. S1.** Partial amino acid sequences encoded by the newly identified avian *CXCL17* gene relics. **(A)** Alignment of partial amino acid sequences encoded by avian *CXCL17* gene relics. XX represents omitted sequence. (**B**) phylogenetic tree of the partial amino acid sequences encoded by avian *CXCL17* gene relics.

#### 6xHis-Hs-Sc-CXCL17

```
1    ATG CAT CAC CAT CAC CAC CAT AGC AGT CTG AAT CCA GGT GTC GCA CGT GGC CAC CGC GAC CGT GGC CAG GCT TCT CGC CGT TGG CTC CAG
     TAC GTA GTG GTA GTG GTG GTA TCG TCA GAC TTA GGT CCA CAG CGT GCA CCG GTG GCG CTG GCA CCG GTC CGA AGA GCG GCA ACC GAG GTC
      M   H   H   H   H   H   H   S   S   L   N   P   G   V   A   R   G   H   R   D   R   G   Q   A   S   R   R   W   L   Q

91   GAA GGT GGC CAA GAA TGT GAG TGC AAA GAT TGG TTC CTG CGT GCC CCG CGC CGT AAA TTC ATG ACC GTG TCC GGT CTG CCG AAG AAA CAG
     CTT CCA CCG GTT CTT ACA CTC ACG TTT CTA ACC AAG GAC GCA CGG GGC GCG GCA TTT AAG TAC TGG CAC AGG CCA GAC GGC TTC TTT GTC
      E   G   G   Q   E   C   E   C   K   D   W   F   L   R   A   P   R   R   K   F   M   T   V   S   G   L   P   K   K   Q

181  TGC CCT TGT GAT CAT TTC AAG GGC AAT GTT AAG AAA ACG CGC CAC CAA CGT CAT CAC CGT AAG CCG AAC AAA CAT TCA CGC CGC TGC CCG
     ACG GGA ACA CTA GTA AAG TTC CCG TTA CAA TTC TTT TGC GCG GTG GTT GCA GTA GTG GCA TTC GGC TTG TTT GTA AGT GCG GCG ACG GGC
      C   P   C   D   H   F   K   G   N   V   K   K   T   R   H   Q   R   H   H   R   K   P   N   K   H   S   R   R   C   P

271  CCA CCT GCT CGT CGT TGT CGC CCG CGT GCA CCA ACC AAA GGT CTG TCT TTA CCG CTG TAA
     GGT GGA CGA GCA GCA ACA GCG GGC GCA CGT GGT TGG TTT CCA GAC AGA AAT GGC GAC ATT
      P   P   A   R   R   C   R   P   R   A   P   T   K   G   L   S   L   P   L   *
```

#### 6xHis-Cc-Sc-CXCL17

```
1    ATG CAT CAT CAC CAC CAT CAC AGC CCG CAC GTG CCA GGT GGC GGT CAG GGC GCG GAA GGT GAG CTG AAC CCT ACC CGT CGC CGT GCA ACG
     TAC GTA GTA GTG GTG GTA GTG TCG GGC GTG CAC GGT CCA CCG CCA GTC CCG CGC CTT CCA CTC GAC TTG GGA TGG GCA GCG GCA CGT TGC
      M   H   H   H   H   H   H   S   P   H   V   P   G   G   G   Q   G   A   E   G   E   L   N   P   T   R   R   R   A   T

91   GGC GCC GGC GTG TGT GAA TGC CCG GGC TGG AGT GGT TCT CGT CGC CAA CTG GCT GGC GAT CTC CGT GTT GGT GCA CGT CAC AAT GGC CTG
     CCG CGG CCG CAC ACA CTT ACG GGC CCG ACC TCA CCA AGA GCA GCG GTT GAC CGA CCG CTA GAG GCA CAA CCA CGT GCA GTG TTA CCG GAC
      G   A   G   V   C   E   C   P   G   W   S   G   S   R   R   Q   L   A   G   D   L   R   V   G   A   R   H   N   G   L

181  TGT CCA TGC GAC TCC ATT GTA CCT CAC GGC CAG GGT CGT GGC CGC CAT CCG CAG GGT CGT GGC CGT CGC GGC CAA CAT CAC GGC CGT AAA
     ACA GGT ACG CTG AGG TAA CAT GGA GTG CCG GTC CCA GCA CCG GCG GTA GGC GTC CCA GCA CCG GCA GCG CCG GTT GTA GTG CCG GCA TTT
      C   P   C   D   S   I   V   P   H   G   Q   G   R   G   R   H   P   Q   G   R   G   R   R   G   Q   H   H   G   R   K

271  CAG CAA CGC TGC CCG CCA CCT GCT CGT CGT TGT CGC CCG CGT GCA CCA ACC AAA GGT CTG TCT TTA CCG CTG TAA
     GTC GTT GCG ACG GGC GGT GGA CGA GCA GCA ACA GCG GGC GCA CGT GGT TGG TTT CCA GAC AGA AAT GGC GAC ATT
      Q   Q   R   C   P   P   P   A   R   R   C   R   P   R   A   P   T   K   G   L   S   L   P   L   *
```

#### 6xHis-SmBiT-Cc-Sc-CXCL17

```
1    ATG CAT CAT CAC CAC CAT CAC GGT GTG ACC GGC TAC CGT CTG TTT GAA GAA ATT CTG GGC GGC AGC CCG CAC GTG CCA GGT GGC GGT CAG
     TAC GTA GTA GTG GTG GTA GTG CCA CAC TGG CCG ATG GCA GAC AAA CTT CTT TAA GAC CCG CCG TCG GGC GTG CAC GGT CCA CCG CCA GTC
      M   H   H   H   H   H   H   G   V   T   G   Y   R   L   F   E   E   I   L   G   G   S   P   H   V   P   G   G   G   Q

91   GGC GCG GAA GGT GAG CTG AAC CCT ACC CGT CGC CGT GCA ACG GGC GCC GGC GTG TGT GAA TGC CCG GGC TGG AGT GGT TCT CGT CGC CAA
     CCG CGC CTT CCA CTC GAC TTG GGA TGG GCA GCG GCA CGT TGC CCG CGG CCG CAC ACA CTT ACG GGC CCG ACC TCA CCA AGA GCA GCG GTT
      G   A   E   G   E   L   N   P   T   R   R   R   A   T   G   A   G   V   C   E   C   P   G   W   S   G   S   R   R   Q

181  CTG GCT GGC GAT CTC CGT GTT GGT GCA CGT CAC AAT GGC CTG TGT CCA TGC GAC TCC ATT GTA CCT CAC GGC CAG GGT CGT GGC CGC CAT
     GAC CGA CCG CTA GAG GCA CAA CCA CGT GCA GTG TTA CCG GAC ACA GGT ACG CTG AGG TAA CAT GGA GTG CCG GTC CCA GCA CCG GCG GTA
      L   A   G   D   L   R   V   G   A   R   H   N   G   L   C   P   C   D   S   I   V   P   H   G   Q   G   R   G   R   H

271  CCG CAG GGT CGT GGC CGT CGC GGC CAA CAT CAC GGC CGT AAA CAG CAA CGC TGC CCG CCA CCT GCT CGT CGT TGT CGC CCG CGT GCA CCA
     GGC GTC CCA GCA CCG GCA GCG CCG GTT GTA GTG CCG GCA TTT GTC GTT GCG ACG GGC GGT GGA CGA GCA GCA ACA GCG GGC GCA CGT GGT
      P   Q   G   R   G   R   R   G   Q   H   H   G   R   K   Q   Q   R   C   P   P   P   A   R   R   C   R   P   R   A   P

361  ACC AAA GGT CTG TCT TTA CCG CTG TAA
     TGG TTT CCA GAC AGA AAT GGC GAC ATT
      T   K   G   L   S   L   P   L   *
```

**Fig. S2.** The nucleotide sequence and amino acid sequence of hybrid CXCL17s carrying a C-terminal fragment encoded by African ostrich *CXCL17* gene relic. The amino acid sequence of Cc-CXCL17 or Hs-CXCL17 is shown in green, that encoded by the African ostrich gene relic is shown in red, the SmBiT-tag is shown in blue, and the 6×His-tag in black.

### Sc-GPR25 in pEAZY vector

```
1     GGC GCT GGG GAT GCC GAC ATG GGT CCC CAA GCA GCC GAG GAC TTC GTG CTG TCC GCA GAC TAC GAT TAC CCA GTG GGG ACC AAA TCC ACG
      CCG CGA CCC CTA CGG CTG TAC CCA GGG GTT CGT CGG CTC CTG AAG CAC GAC AGG CGT CTG ATG CTA ATG GGT CAC CCC TGG TTT AGG TGC
                               M   G   P   Q   A   A   E   D   F   V   L   S   A   D   Y   D   Y   P   V   G   T   K   S   T

91    GGG ACG GAG GAG CGT TTC TCC AGG TGG GAG TCG TCC TAC ATG ACC GTC TTC ATC CCC ATC CTC TAC TCC TTC ATT TTC CTC CTT GGC TTC
      CCC TGC CTC CTC GCA AAG AGG TCC ACC CTC AGC AGG ATG TAC TGG CAG AAG TAG GGG TAG GAG ATG AGG AAG TAA AAG GAG GAA CCG AAG
       G   T   E   E   R   F   S   R   W   E   S   S   Y   M   T   V   F   I   P   I   L   Y   S   F   I   F   L   L   G   F

181   GTG GGG AAC CTC TTT GTG ATC GTG CTG CTG GCC AAG AAG AGA GGC AGC AAG AGG ATG GTG GAT ACG TTT GTG CTG AAC CTG GCT GTG GCC
      CAC CCC TTG GAG AAA CAC TAG CAC GAC GAC CGG TTC TTC TCT CCG TCG TTC TCC TAC CAC CTA TGC AAA CAC GAC TTG GAC CGA CAC CGG
       V   G   N   L   F   V   I   V   L   L   A   K   K   R   G   S   K   R   M   V   D   T   F   V   L   N   L   A   V   A

271   GAC GTG ATC TTC GTC TGC ACC CTG CCC TTC TGG GTC GTG GCA GGC GCC CAT GGG AAC CGC TGG CAG CTC GGG GAA GGG CTC TGC AAG GTG
      CTG CAC TAG AAG CAG ACG TGG GAC GGG AAG ACC CAG CAC CGT CCG CGG GTA CCC TTG GCG ACC GTC GAG CCC CTT CCC GAG ACG TTC CAC
       D   V   I   F   V   C   T   L   P   F   W   V   V   A   G   A   H   G   N   R   W   Q   L   G   E   G   L   C   K   V

361   AGC AGC TAC GCC ATC ACT GTG AAC CGC TGC TCC AGC ATC CTC TTC CTC ACC GCC CTG AGC GTG GAG CGC TAC CTG GTC ATC AGG AAG GTG
      TCG TCG ATG CGG TAG TGA CAC TTG GCG ACG AGG TCG TAG GAG AAG GAG TGG CGG GAC TCG CAC CTC GCG ATG GAC CAG TAG TCC TTC CAC
       S   S   Y   A   I   T   V   N   R   C   S   S   I   L   F   L   T   A   L   S   V   E   R   Y   L   V   I   R   K   V

451   CTG GAT ACC AAG ATG ACG GGG TCC AAG AGG CAC GTC TTT GTC ACT TGT GGG CTG ATC TGG GTT GTG TCC CTC CTC CTG GGT GCG CCG TCC
      GAC CTA TGG TTC TAC TGC CCC AGG TTC TCC GTG CAG AAA CAG TGA ACA CCC GAC TAG ACC CAA CAC AGG GAG GAG GAC CCA CGC GGC AGG
       L   D   T   K   M   T   G   S   K   R   H   V   F   V   T   C   G   L   I   W   V   V   S   L   L   L   G   A   P   S

541   CTG GAG TAC CGG CGG CTA GAT GGA GAT GAC TGT TGG GAT GAG GAT GGG GAG GAT TTC AGC CTG GCA ATG GTC TTC CTC ACC TTC CTC CTG
      GAC CTC ATG GCC GCC GAT CTA CCT CTA CTG ACA ACC CTA CTC CTA CCC CTC CTA AAG TCG GAC CGT TAC CAG AAG GAG TGG AAG GAG GAC
       L   E   Y   R   R   L   D   G   D   D   C   W   D   E   D   G   E   D   F   S   L   A   M   V   F   L   T   F   L   L

631   CCA CTG GCG GTC ATC TTG TTC TGC TAC TGC TCC ATC TAC TAC CAG CTC CAC AGG CAC ATC AGG CTG GGG AAG GGG ATG AGG AAC TCC CAC
      GGT GAC CGC CAG TAG AAC AAG ACG ATG ACG AGG TAG ATG ATG GTC GAG GTG TCC GTG TAG TCC GAC CCC TTC CCC TAC TCC TTG AGG GTG
       P   L   A   V   I   L   F   C   Y   C   S   I   Y   Y   Q   L   H   R   H   I   R   L   G   K   G   M   R   N   S   H

721   CGG ATC ATC TTC ACC ATC GTC GGA GCC TTC ATC TGC TCC TGG CTG CCC TTG AAT GTC TGC AAG GTG TTG CTC TTC TTC ATC GCG AAG GGG
      GCC TAG TAG AAG TGG TAG CAG CCT CGG AAG TAG ACG AGG ACC GAC GGG AAC TTA CAG ACG TTC CAC AAC GAG AAG AAG TAG CGC TTC CCC
       R   I   I   F   T   I   V   G   A   F   I   C   S   W   L   P   L   N   V   C   K   V   L   L   F   F   I   A   K   G

811   ATG CTG GTC CTC TCC GAG GTG GAG GAG GAG GCC TTG AGG TGG GTC GTC ACC AGC AGC ACA TGC CTG GCC TTC ATG AAC AGC TGC GTC AAC
      TAC GAC CAG GAG AGG CTC CAC CTC CTC CTC CGG AAC TCC ACC CAG CAG TGG TCG TCG TGT ACG GAC CGG AAG TAC TTG TCG ACG CAG TTG
       M   L   V   L   S   E   V   E   E   E   A   L   R   W   V   V   T   S   S   T   C   L   A   F   M   N   S   C   V   N

901   CCC ATC ATT TAC ACC CTG ATG GAT CGC CAT TTC CGG CGC CAG CTG CTG CCC CCG GCT CTC CGG GTG CTC TGC CTT TCC CCC GCA GAG AAG
      GGG TAG TAA ATG TGG GAC TAC CTA GCG GTA AAG GCC GCG GTC GAC GAC GGG GGC CGA GAG GCC CAC GAG ACG GAA AGG GGG CGT CTC TTC
       P   I   I   Y   T   L   M   D   R   H   F   R   R   Q   L   L   P   P   A   L   R   V   L   C   L   S   P   A   E   K

991   GGG CAC AAC ATG GCC ACG ACG TCC TCT GTG ACC GAG TCC AGC CTC ATG TTC GGC GTC AGG ACC AAG CCG TCC GCA CCG CCG GGG CGC TGG
      CCC GTG TTG TAC CGG TGC TGC AGG AGA CAC TGG CTC AGG TCG GAG TAC AAG CCG CAG TCC TGG TTC GGC AGG CGT GGC GGC CCC GCG ACC
       G   H   N   M   A   T   T   S   S   V   T   E   S   S   L   M   F   G   V   R   T   K   P   S   A   P   P   G   R   W

1081  CCA GCA GTG CAA GAG GAG AAG GAT TGG GGG CCA AAC GAG TAG GTC ATT TGG GGA GGG CTG AGC TTG
      GGT CGT CAC GTT CTC CTC TTC CTA ACC CCC GGT TTG CTC ATC CAG TAA ACC CCT CCC GAC TCG AAC
       P   A   V   Q   E   E   K   D   W   G   P   N   E   *
```

### Sc-GPR25-LgBiT in pTRE3G-BI (coexpressing with SmBiT-ARRB2 via a bi-directional promoter)

```
1     CGG GGA GAC CCA AGC TGG ATG GGT CCC CAA GCA GCC GAG GAC TTC GTG CTG TCC GCA GAC TAC GAT TAC CCA GTG GGG ACC AAA TCC ACG
      GCC CCT CTG GGT TCG ACC TAC CCA GGG GTT CGT CGG CTC CTG AAG CAC GAC AGG CGT CTG ATG CTA ATG GGT CAC CCC TGG TTT AGG TGC
                               M   G   P   Q   A   A   E   D   F   V   L   S   A   D   Y   D   Y   P   V   G   T   K   S   T

91    GGG ACG GAG GAG CGT TTC TCC AGG TGG GAG TCG TCC TAC ATG ACC GTC TTC ATC CCC ATC CTC TAC TCC TTC ATT TTC CTC CTT GGC TTC
      CCC TGC CTC CTC GCA AAG AGG TCC ACC CTC AGC AGG ATG TAC TGG CAG AAG TAG GGG TAG GAG ATG AGG AAG TAA AAG GAG GAA CCG AAG
       G   T   E   E   R   F   S   R   W   E   S   S   Y   M   T   V   F   I   P   I   L   Y   S   F   I   F   L   L   G   F

181   GTG GGG AAC CTC TTT GTG ATC GTG CTG CTG GCC AAG AAG AGA GGC AGC AAG AGG ATG GTG GAT ACG TTT GTG CTG AAC CTG GCT GTG GCC
      CAC CCC TTG GAG AAA CAC TAG CAC GAC GAC CGG TTC TTC TCT CCG TCG TTC TCC TAC CAC CTA TGC AAA CAC GAC TTG GAC CGA CAC CGG
       V   G   N   L   F   V   I   V   L   L   A   K   K   R   G   S   K   R   M   V   D   T   F   V   L   N   L   A   V   A

271   GAC GTG ATC TTC GTC TGC ACC CTG CCC TTC TGG GTC GTG GCA GGC GCC CAT GGG AAC CGC TGG CAG CTC GGG GAA GGG CTC TGC AAG GTG
      CTG CAC TAG AAG CAG ACG TGG GAC GGG AAG ACC CAG CAC CGT CCG CGG GTA CCC TTG GCG ACC GTC GAG CCC CTT CCC GAG ACG TTC CAC
       D   V   I   F   V   C   T   L   P   F   W   V   V   A   G   A   H   G   N   R   W   Q   L   G   E   G   L   C   K   V

361   AGC AGC TAC GCC ATC ACT GTG AAC CGC TGC TCC AGC ATC CTC TTC CTC ACC GCC CTG AGC GTG GAG CGC TAC CTG GTC ATC AGG AAG GTG
      TCG TCG ATG CGG TAG TGA CAC TTG GCG ACG AGG TCG TAG GAG AAG GAG TGG CGG GAC TCG CAC CTC GCG ATG GAC CAG TAG TCC TTC CAC
       S   S   Y   A   I   T   V   N   R   C   S   S   I   L   F   L   T   A   L   S   V   E   R   Y   L   V   I   R   K   V

451   CTG GAT ACC AAG ATG ACG GGG TCC AAG AGG CAC GTC TTT GTC ACT TGT GGG CTG ATC TGG GTT GTG TCC CTC CTC CTG GGT GCG CCG TCC
      GAC CTA TGG TTC TAC TGC CCC AGG TTC TCC GTG CAG AAA CAG TGA ACA CCC GAC TAG ACC CAA CAC AGG GAG GAG GAC CCA CGC GGC AGG
       L   D   T   K   M   T   G   S   K   R   H   V   F   V   T   C   G   L   I   W   V   V   S   L   L   L   G   A   P   S

541   CTG GAG TAC CGG CGG CTA GAT GGA GAT GAC TGT TGG GAT GAG GAT GGG GAG GAT TTC AGC CTG GCA ATG GTC TTC CTC ACC TTC CTC CTG
      GAC CTC ATG GCC GCC GAT CTA CCT CTA CTG ACA ACC CTA CTC CTA CCC CTC CTA AAG TCG GAC CGT TAC CAG AAG GAG TGG AAG GAG GAC
       L   E   Y   R   R   L   D   G   D   D   C   W   D   E   D   G   E   D   F   S   L   A   M   V   F   L   T   F   L   L

631   CCA CTG GCG GTC ATC TTG TTC TGC TAC TGC TCC ATC TAC TAC CAG CTC CAC AGG CAC ATC AGG CTG GGG AAG GGG ATG AGG AAC TCC CAC
      GGT GAC CGC CAG TAG AAC AAG ACG ATG ACG AGG TAG ATG ATG GTC GAG GTG TCC GTG TAG TCC GAC CCC TTC CCC TAC TCC TTG AGG GTG
       P   L   A   V   I   L   F   C   Y   C   S   I   Y   Y   Q   L   H   R   H   I   R   L   G   K   G   M   R   N   S   H

721   CGG ATC ATC TTC ACC ATC GTC GGA GCC TTC ATC TGC TCC TGG CTG CCC TTG AAT GTC TGC AAG GTG TTG CTC TTC TTC ATC GCG AAG GGG
```

GCC TAG TAG AAG TGG TAG CAG CCT CGG AAG TAG ACG AGG ACC GAC GGG AAC TTA CAG ACG TTC CAC AAC GAG AAG AAG TAG CGC TTC CCC
R I I F T I V G A F I C S W L P L N V C K V L L F F I A K G

811 ATG CTG GTC CTC TCC GAG GTG GAG GAG GAG GCC TTG AGG TGG GTC GTC ACC AGC AGC ACA TGC CTG GCC TTC ATG AAC AGC TGC GTC AAC
TAC GAC CAG GAG AGG CTC CAC CTC CTC CTC CGG AAC TCC ACC CAG CAG TGG TCG TCG TGT ACG GAC CGG AAG TAC TTG TCG ACG CAG TTG
M L V L S E V E E E A L R W V V T S S T C L A F M N S C V N

901 CCC ATC ATT TAC ACC CTG ATG GAT CGC CAT TTC CGG CGC CAG CTG CTG CCC CCG GCT CTC CGG GTG CTC TGC CTT TCC CCC GCA GAG AAG
GGG TAG TAA ATG TGG GAC TAC CTA GCG GTA AAG GCC GCG GTC GAC GAC GGG GGC CGA GAG GCC CAC GAG ACG GAA AGG GGG CGT CTC TTC
P I I Y T L M D R H F R R Q L L P P A L R V L C L S P A E K

991 GGG CAC AAC ATG GCC ACG ACG TCC TCT GTG ACC GAG TCC AGC CTC ATG TTC GGC GTC AGG ACC AAG CCG TCC GCA CCG CCG GGG CGC TGG
CCC GTG TTG TAC CGG TGC TGC AGG AGA CAC TGG CTC AGG TCG GAG TAC AAG CCG CAG TCC TGG TTC GGC AGG CGT GGC GGC CCC GCG ACC
G H N M A T T S S V T E S S L M F G V R T K P S A P P G R W

1081 CCA GCA GTG CAA GAG GAG AAG GAT TGG GGG CCA AAC GAG CCA CCG GTC GGT ACC GGC GGA GGG TCT AGC AGT GGC GGT GGG ATG GTC TTC
GGT CGT CAC GTT CTC CTC TTC CTA ACC CCC GGT TTG CTC GGT GGC CAG CCA TGG CCG CCT CCC AGA TCG TCA CCG CCA CCC TAC CAG AAG
P A V Q E E K D W G P N E P P V G T G G G S S S G G G M V F

1171 ACA CTC GAA GAT TTC GTT GGG GAC TGG GAA CAG ACA GCC GCC TAC AAC CTG GAC CAA GTC CTT GAA CAG GGA GGT GTG TCC AGT TTG CTG
TGT GAG CTT CTA AAG CAA CCC CTG ACC CTT GTC TGT CGG CGG ATG TTG GAC CTG GTT CAG GAA CTT GTC CCT CCA CAC AGG TCA AAC GAC
T L E D F V G D W E Q T A A Y N L D Q V L E Q G G V S S L L

1261 CAG AAT CTC GCC GTG TCC GTA ACT CCG ATC CAA AGG ATT GTC CGG AGC GGT GAA AAT GCC CTG AAG ATC GAC ATC CAT GTC ATC ATC CCG
GTC TTA GAG CGG CAC AGG CAT TGA GGC TAG GTT TCC TAA CAG GCC TCG CCA CTT TTA CGG GAC TTC TAG CTG TAG GTA CAG TAG TAG GGC
Q N L A V S V T P I Q R I V R S G E N A L K I D I H V I I P

1351 TAT GAA GGT CTG AGC GCC GAC CAA ATG GCC CAG ATC GAA GAG GTG TTT AAG GTG GTG TAC CCT GTG GAT GAT CAT CAC TTT AAG GTG ATC
ATA CTT CCA GAC TCG CGG CTG GTT TAC CGG GTC TAG CTT CTC CAC AAA TTC CAC CAC ATG GGA CAC CTA CTA GTA GTG AAA TTC CAC TAG
Y E G L S A D Q M A Q I E E V F K V V Y P V D D H H F K V I

1441 CTG CCC TAT GGC ACA CTG GTA ATC GAC GGG GTT ACG CCG AAC ATG CTG AAC TAT TTC GGA CGG CCG TAT GAA GGC ATC GCC GTG TTC GAC
GAC GGG ATA CCG TGT GAC CAT TAG CTG CCC CAA TGC GGC TTG TAC GAC TTG ATA AAG CCT GCC GGC ATA CTT CCG TAG CGG CAC AAG CTG
L P Y G T L V I D G V T P N M L N Y F G R P Y E G I A V F D

1531 GGC AAA AAG ATC ACT GTA ACA GGG ACC CTG TGG AAC GGC AAC AAA ATT ATC GAC GAG CGC CTG ATC ACC CCC GAC GGC TCC ATG CTG TTC
CCG TTT TTC TAG TGA CAT TGT CCC TGG GAC ACC TTG CCG TTG TTT TAA TAG CTG CTC GCG GAC TAG TGG GGG CTG CCG AGG TAC GAC AAG
G K K I T V T G T L W N G N K I I D E R L I T P D G S M L F

1621 CGA GTA ACC ATC AAC AGT TAA
GCT CAT TGG TAG TTG TCA ATT
R V T I N S *

### sLgBiT-Sc-GPR25 in PB-TRE vector

1 ATG AAC TCC TTC TCC ACA AGC GCC TTC GGT CCA GTT GCC TTC TCC CTG GGC CTG CTC CTG GTG TTG CCT GCT GCC TTC CCT GCC CCA GTC
TAC TTG AGG AAG AGG TGT TCG CGG AAG CCA GGT CAA CGG AAG AGG GAC CCG GAC GAG GAC CAC AAC GGA CGA CGG AAG GGA CGG GGT CAG
M N S F S T S A F G P V A F S L G L L L V L P A A F P A P V

91 TTC ACA CTC GAA GAT TTC GTT GGG GAC TGG GAA CAG ACA GCC GCC TAC AAC CTG GAC CAA GTC CTT GAA CAG GGA GGT GTG TCC AGT TTG
AAG TGT GAG CTT CTA AAG CAA CCC CTG ACC CTT GTC TGT CGG CGG ATG TTG GAC CTG GTT CAG GAA CTT GTC CCT CCA CAC AGG TCA AAC
F T L E D F V G D W E Q T A A Y N L D Q V L E Q G G V S S L

181 CTG CAG AAT CTC GCC GTG TCC GTA ACT CCG ATC CAA AGG ATT GTC CGG AGC GGT GAA AAT GCC CTG AAG ATC GAC ATC CAT GTC ATC ATC
GAC GTC TTA GAG CGG CAC AGG CAT TGA GGC TAG GTT TCC TAA CAG GCC TCG CCA CTT TTA CGG GAC TTC TAG CTG TAG GTA CAG TAG TAG
L Q N L A V S V T P I Q R I V R S G E N A L K I D I H V I I

271 CCG TAT GAA GGT CTG AGC GCC GAC CAA ATG GCC CAG ATC GAA GAG GTG TTT AAG GTG GTG TAC CCT GTG GAT GAT CAT CAC TTT AAG GTG
GGC ATA CTT CCA GAC TCG CGG CTG GTT TAC CGG GTC TAG CTT CTC CAC AAA TTC CAC CAC ATG GGA CAC CTA CTA GTA GTG AAA TTC CAC
P Y E G L S A D Q M A Q I E E V F K V V Y P V D D H H F K V

361 ATC CTG CCC TAT GGC ACA CTG GTA ATC GAC GGG GTT ACG CCG AAC ATG CTG AAC TAT TTC GGA CGG CCG TAT GAA GGC ATC GCC GTG TTC
TAG GAC GGG ATA CCG TGT GAC CAT TAG CTG CCC CAA TGC GGC TTG TAC GAC TTG ATA AAG CCT GCC GGC ATA CTT CCG TAG CGG CAC AAG
I L P Y G T L V I D G V T P N M L N Y F G R P Y E G I A V F

451 GAC GGC AAA AAG ATC ACT GTA ACA GGG ACC CTG TGG AAC GGC AAC AAA ATT ATC GAC GAG CGC CTG ATC ACC CCC GAC GGC TCC ATG CTG
CTG CCG TTT TTC TAG TGA CAT TGT CCC TGG GAC ACC TTG CCG TTG TTT TAA TAG CTG CTC GCG GAC TAG TGG GGG CTG CCG AGG TAC GAC
D G K K I T V T G T L W N G N K I I D E R L I T P D G S M L

541 TTC CGA GTA ACC ATC AAC AGT GGT GGC GGC TCT GGT GGT GGC AGC GGC GGT GGT ACC GGT CCC CAA GCA GCC GAG GAC TTC GTG CTG TCC
AAG GCT CAT TGG TAG TTG TCA CCA CCG CCG AGA CCA CCA CCG TCG CCG CCA CCA TGG CCA GGG GTT CGT CGG CTC CTG AAG CAC GAC AGG
F R V T I N S G G G S G G G S G G G T G P Q A A E D F V L S

631 GCA GAC TAC GAT TAC CCA GTG GGG ACC AAA TCC ACG GGG ACG GAG GAG CGT TTC TCC AGG TGG GAG TCG TCC TAC ATG ACC GTC TTC ATC
CGT CTG ATG CTA ATG GGT CAC CCC TGG TTT AGG TGC CCC TGC CTC CTC GCA AAG AGG TCC ACC CTC AGC AGG ATG TAC TGG CAG AAG TAG
A D Y D Y P V G T K S T G T E E R F S R W E S S Y M T V F I

721 CCC ATC CTC TAC TCC TTC ATT TTC CTC CTT GGC TTC GTG GGG AAC CTC TTT GTG ATC GTG CTG CTG GCC AAG AAG AGA GGC AGC AAG AGG
GGG TAG GAG ATG AGG AAG TAA AAG GAG GAA CCG AAG CAC CCC TTG GAG AAA CAC TAG CAC GAC GAC CGG TTC TTC TCT CCG TCG TTC TCC
P I L Y S F I F L L G F V G N L F V I V L L A K K R G S K R

811 ATG GTG GAT ACG TTT GTG CTG AAC CTG GCT GTG GCC GAC GTG ATC TTC GTC TGC ACC CTG CCC TTC TGG GTC GTG GCA GGC GCC CAT GGG
TAC CAC CTA TGC AAA CAC GAC TTG GAC CGA CAC CGG CTG CAC TAG AAG CAG ACG TGG GAC GGG AAG ACC CAG CAC CGT CCG CGG GTA CCC
M V D T F V L N L A V A D V I F V C T L P F W V V A G A H G

901 AAC CGC TGG CAG CTC GGG GAA GGG CTC TGC AAG GTG AGC AGC TAC GCC ATC ACT GTG AAC CGC TGC TCC AGC ATC CTC TTC CTC ACC GCC
TTG GCG ACC GTC GAG CCC CTT CCC GAG ACG TTC CAC TCG TCG ATG CGG TAG TGA CAC TTG GCG ACG AGG TCG TAG GAG AAG GAG TGG CGG
N R W Q L G E G L C K V S S Y A I T V N R C S S I L F L T A

```
991   CTG AGC GTG GAG CGC TAC CTG GTC ATC AGG AAG GTG CTG GAT ACC AAG ATG ACG GGG TCC AAG AGG CAC GTC TTT GTC ACT TGT GGG CTG
      GAC TCG CAC CTC GCG ATG GAC CAG TAG TCC TTC CAC GAC CTA TGG TTC TAC TGC CCC AGG TTC TCC GTG CAG AAA CAG TGA ACA CCC GAC
       L   S   V   E   R   Y   L   V   I   R   K   V   L   D   T   K   M   T   G   S   K   R   H   V   F   V   T   C   G   L

1081  ATC TGG GTT GTG TCC CTC CTC CTG GGT GCG CCG TCC CTG GAG TAC CGG CGG CTA GAT GGA GAT GAC TGT TGG GAT GAG GAT GGG GAG GAT
      TAG ACC CAA CAC AGG GAG GAG GAC CCA CGC GGC AGG GAC CTC ATG GCC GCC GAT CTA CCT CTA CTG ACA ACC CTA CTC CTA CCC CTC CTA
       I   W   V   V   S   L   L   L   G   A   P   S   L   E   Y   R   R   L   D   G   D   D   C   W   D   E   D   G   E   D

1171  TTC AGC CTG GCA ATG GTC TTC CTC ACC TTC CTC CTG CCA CTG GCG GTC ATC TTG TTC TGC TAC TGC TCC ATC TAC TAC CAG CTC CAC AGG
      AAG TCG GAC CGT TAC CAG AAG GAG TGG AAG GAG GAC GGT GAC CGC CAG TAG AAC AAG ACG ATG ACG AGG TAG ATG ATG GTC GAG GTG TCC
       F   S   L   A   M   V   F   L   T   F   L   L   P   L   A   V   I   L   F   C   Y   C   S   I   Y   Y   Q   L   H   R

1261  CAC ATC AGG CTG GGG AAG GGG ATG AGG AAC TCC CAC CGG ATC ATC TTC ACC ATC GTC GGA GCC TTC ATC TGC TCC TGG CTG CCC TTG AAT
      GTG TAG TCC GAC CCC TTC CCC TAC TCC TTG AGG GTG GCC TAG TAG AAG TGG TAG CAG CCT CGG AAG TAG ACG AGG ACC GAC GGG AAC TTA
       H   I   R   L   G   K   G   M   R   N   S   H   R   I   I   F   T   I   V   G   A   F   I   C   S   W   L   P   L   N

1351  GTC TGC AAG GTG TTG CTC TTC TTC ATC GCG AAG GGG ATG CTG GTC CTC TCC GAG GTG GAG GAG GAG GCC TTG AGG TGG GTC GTC ACC AGC
      CAG ACG TTC CAC AAC GAG AAG AAG TAG CGC TTC CCC TAC GAC CAG GAG AGG CTC CAC CTC CTC CTC CGG AAC TCC ACC CAG CAG TGG TCG
       V   C   K   V   L   L   F   F   I   A   K   G   M   L   V   L   S   E   V   E   E   E   A   L   R   W   V   V   T   S

1441  AGC ACA TGC CTG GCC TTC ATG AAC AGC TGC GTC AAC CCC ATC ATT TAC ACC CTG ATG GAT CGC CAT TTC CGG CGC CAG CTG CTG CCC CCG
      TCG TGT ACG GAC CGG AAG TAC TTG TCG ACG CAG TTG GGG TAG TAA ATG TGG GAC TAC CTA GCG GTA AAG GCC GCG GTC GAC GAC GGG GGC
       S   T   C   L   A   F   M   N   S   C   V   N   P   I   I   Y   T   L   M   D   R   H   F   R   R   Q   L   L   P   P

1531  GCT CTC CGG GTG CTC TGC CTT TCC CCC GCA GAG AAG GGG CAC AAC ATG GCC ACG ACG TCC TCT GTG ACC GAG TCC AGC CTC ATG TTC GGC
      CGA GAG GCC CAC GAG ACG GAA AGG GGG CGT CTC TTC CCC GTG TTG TAC CGG TGC TGC AGG AGA CAC TGG CTC AGG TCG GAG TAC AAG CCG
       A   L   R   V   L   C   L   S   P   A   E   K   G   H   N   M   A   T   T   S   S   V   T   E   S   S   L   M   F   G

1621  GTC AGG ACC AAG CCG TCC GCA CCG CCG GGG CGC TGG CCA GCA GTG CAA GAG GAG AAG GAT TGG GGG CCA AAC GAG TAG AAA CCC GCT GAT
      CAG TCC TGG TTC GGC AGG CGT GGC GGC CCC GCG ACC GGT CGT CAC GTT CTC CTC TTC CTA ACC CCC GGT TTG CTC ATC TTT GGG CGA CTA
       V   R   T   K   P   S   A   P   P   G   R   W   P   A   V   Q   E   E   K   D   W   G   P   N   E   *

1711  CAG CCT C
      GTC GGA G
```

**Fig. S3.** The nucleotide sequence and amino acid sequence of Sc-GPR25 in various expression constructs. The amino acid sequence of Sc-GPR25 is shown in red, that of LgBiT in blue. The signal peptide in sLgBiT is shaded.

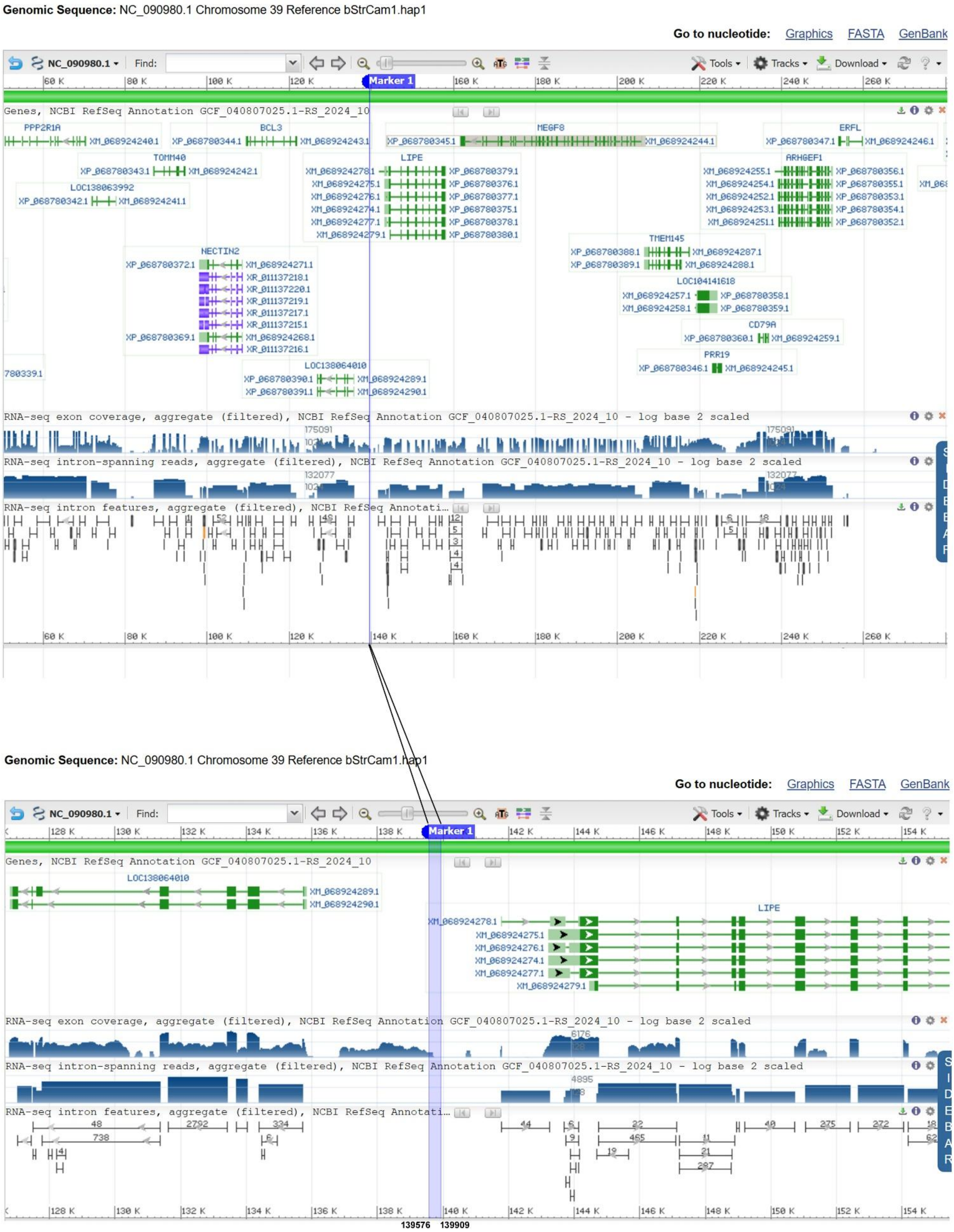


**Fig. S4.** Position of the *CXCL17* gene relic in the reference genome of African ostrich (*Struthio camelus*). The information is downloaded from the NCBI gene database and the *CXCL17* gene relic region (139576–139909) is shown in blue (https://www.ncbi.nlm.nih.gov/gene/?term=lipe%2C+ostrich).

**A**

**African ostrich *CXCL17* gene relic from NCBI database**

```
1      CGG GCT CCG GCC TCC GCC ATG CGC CGC CCG CTC GCC CTG GCC CTG CTG CTG CTG CTG CTG CTG CTG CTG GGG GGG GGG GAC GGG ACC CCC
       GCC CGA GGC CGG AGG CGG TAC GCG GCG GGC GAG CGG GAC CGG GAC GAC GAC GAC GAC GAC GAC GAC GAC CCC CCC CCC CTG CCC TGG GGG
                                M   R   R   P   L   A   L   A   L   L   L   L   L   L   L   L   G   G   G   D   G   T   P

91     CGG CCC CCC CGG CCC CCC CGG GGG GCA CCG GCC CCC GTC GGC GCC GGG AGC CGC CGG CCG CTC TGC ACC TGC GGG AGA GGA GCC GCG A
       GCC GGG GGG GCC GGG GGG GCC CCC CGT GGC CGG GGG CAG CCG CGG CCC TCG GCG GCC GGC GAG ACG TGG ACG CCC TCT CCT CGG CGC T

179    GTG AGG CAG CGC CCT TTT TTG CGG GGG GGG GGA TTT TTT GGG GTA CCC CCC CTT TAT TTC TTT CTC TCC CCC CCC CCC GCA GGG GTC CCC
       CAC TCC GTC GCG GGA AAA AAC GCC CCC CCC CCT AAA AAA CCC CAT GGG GGG GAA ATA AAG AAA GAG AGG GGG GGG GGG CGT CCC CAG GGG

269    GCC GCC CCC CGG GGC CGG TGC CCG CCG CCC GCC CGG CGC TGC CGC CCC CGC GCC CCC ACC AAG GGG CTC TCG CTG CCC CTC TAG GGG CCG
       CGG CGG GGG GCC CCG GCC ACG GGC GGC GGG CGG GCC GCG ACG GCG GGG GCG CGG GGG TGG TTC CCC GAG AGC GAC GGG GAG ATC CCC GGC
        A   A   P   R   G   R   C   P   P   P   A   R   R   C   R   P   R   A   P   T   K   G   L   S   L   P   L   *

359    AAT TCG GAT GCC CCC
       TTA AGC CTA CGG GGG
```

**B**

**African ostrich *CXCL17* gene relic cloned from genomic DNA in this study**

```
1      CGG GCT CCG GCC TCC GCC ATG CGC CGC CCG CTC GCC CTG GCC CTG CTG CTG CTG CTG CTG CTG CTG GGG GGG GGG GAC GGG ACC CCC CGG
       GCC CGA GGC CGG AGG CGG TAC GCG GCG GGC GAG CGG GAC CGG GAC GAC GAC GAC GAC GAC GAC GAC CCC CCC CCC CTG CCC TGG GGG GCC
                                M   R   R   P   L   A   L   A   L   L   L   L   L   L   L   L   G   G   G   D   G   T   P   R

91     CCC CCC CGG CCC CCC CGG GGG GCA CCG GCC CCC GTC GGC GCC GGG AGC CGC CGG CCG CTC TGC ACC TGC GGG AGA GGA GCC GCG AGT G
       GGG GGG GCC GGG GGG GCC CCC CGT GGC CGG GGG CAG CCG CGG CCC TCG GCG GCC GGC GAG ACG TGG ACG CCC TCT CCT CGG CGC TCA C

179    AGG CAG CGC CCT TTC TTG CGG GGG GGG GGG ATT TTT TGG GGT ACC CCC CCC CTT TAT TTC TTT CTC TCC CCC CCC CCC GCA GGG GTC CCC
       TCC GTC GCG GGA AAG AAC GCC CCC CCC CCC TAA AAA ACC CCA TGG GGG GGG GAA ATA AAG AAA GAG AGG GGG GGG GGG CGT CCC CAG GGG

269    GCC GCC CCC CGG GGC CGG TGC CCG CCG CCC GCC CGG CGC TGC CGC CCC CGC GCC CCC ACC AAG GGG CTC TCG CTG CCC CTC TAG GGG CCG
       CGG CGG GGG GCC CCG GCC ACG GGC GGC GGG CGG GCC GCG ACG GCG GGG GCG CGG GGG TGG TTC CCC GAG AGC GAC GGG GAG ATC CCC GGC
        A   A   P   R   G   R   C   P   P   P   A   R   R   C   R   P   R   A   P   T   K   G   L   S   L   P   L   *

359    AAT TCG GAT GCC CCC
       TTA AGC CTA CGG GGG
```

**C**

**DNA sequence alignment of the reference genomic sequence and PCR product of the ostrich *CXCL17* gene relic**

```
Reference sequence   (1)  CGGGCTCCGGCCTCCGCCATGCGCCGCCCGCTCGCCCTGGCCCTGCTGCTGCTGCTGCTGCTGCTGGGGGGGGGGACGGGACCCCCCGGCCCCCCC
PCR product          (1)  CGGGCTCCGGCCTCCGCCATGCGCCGCCCGCTCGCCCTGGCCCTGCTGCTGCTGCTGCTGCTG---GGGGGGGGGGACGGGACCCCCCGGCCCCCCC

Reference sequence (101)  GGCCCCCCCGGGGGGCACCGGCCCCCGTCGGCGCCGGGAGCCGCCGGCCGCTCTGCACCTGCGGGAGAGGAGCCGCGAGTGAGGCAGCGCCCTTTTTTGC
PCR product         (98)  GGCCCCCCCGGGGGGCACCGGCCCCCGTCGGCGCCGGGAGCCGCCGGCCGCTCTGCACCTGCGGGAGAGGAGCCGCGAGTGAGGCAGCGCCCTTTCTTGC

Reference sequence (201)  GGGGGGGGGG-ATTTTTTGGGGTACCCCCCC--TTTATTTCTTTCTCTCCCCCCCCCCCGCAGGGGTCCCCGCCGCCCCCCGGGGCCGGTGCCCGCCGCC
PCR product        (198)  GGGGGGGGGGGATTTTTTGGGGTACCCCCCCCCTTTATTTCTTTCTCTCCCCCCCCCCCGCAGGGGTCCCCGCCGCCCCCCGGGGCCGGTGCCCGCCGCC

Reference sequence (298)  CGCCCGGCGCTGCCGCCCCCGCGCCCCCACCAAGGGGCTCTCGCTGCCCCTCTAGGGGCCGAATTCGGATGCCCCC
PCR product        (298)  CGCCCGGCGCTGCCGCCCCCGCGCCCCCACCAAGGGGCTCTCGCTGCCCCTCTAGGGGCCGAATTCGGATGCCCCC
```

**Fig. S5.** Confirming presence of *CXCL17* gene relic in African ostrich. This genomic DNA sequence is near to the *LIPE* gene and encodes an N-terminal signal peptide (shaded) by its 3’-region and a C-terminal sequence with characteristic features of CXCL17 orthologs by its 3’-region (red), but it no longer encodes an intact CXCL17 protein due to presence of frame-shift mutations. The primer positions for PCR amplification are underlined in panel A and B.

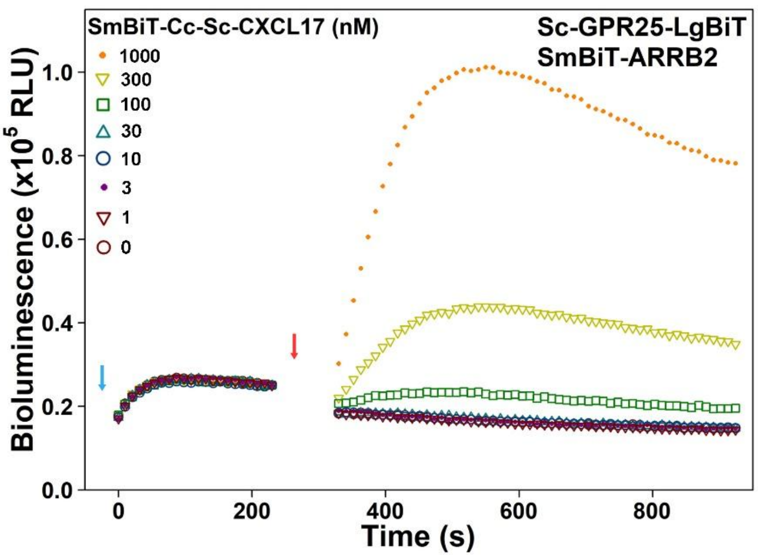


**Fig. S6.** Activity of SmBiT-Cc-Sc-CXCL17 measured via the NanoBiT-based β-arrestin recruitment assay.

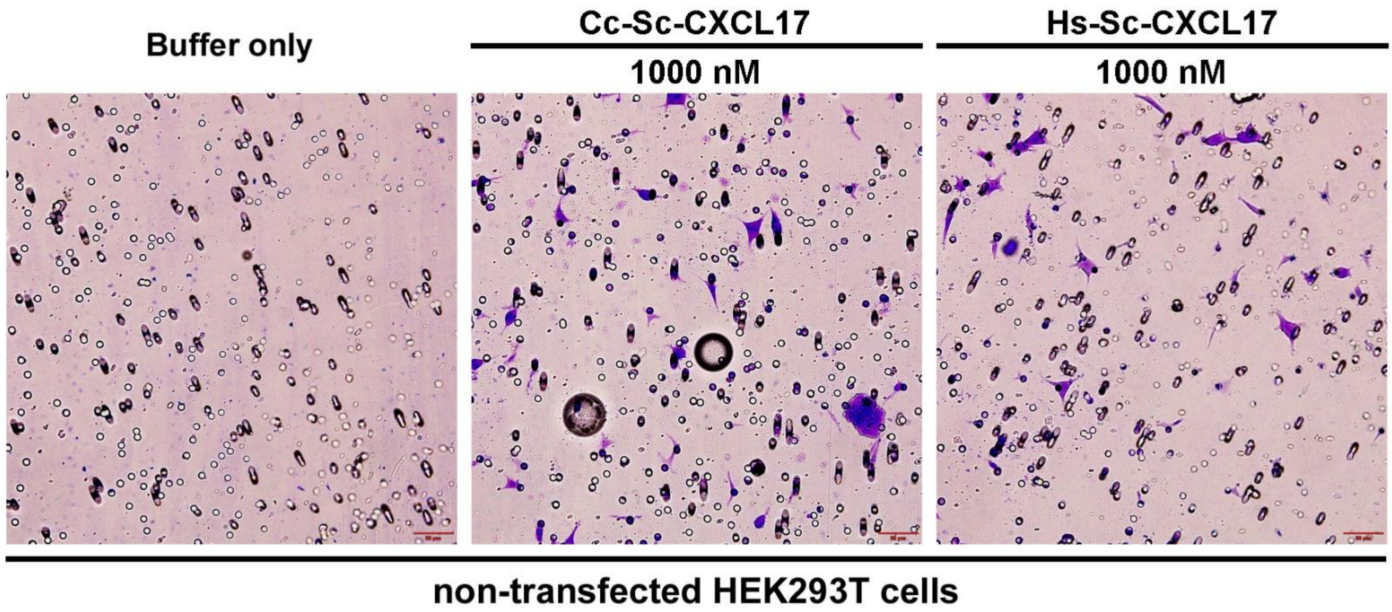


**Fig. S7.** Presentative images of migrated non-transfected HEK293T cells after induced by Cc-Sc-CXCL17 or Hs-Sc-CXCL17 in chemotaxis assays.
